# Transcriptome-wide RNA structure probing with temporal resolution

**DOI:** 10.1101/2023.09.28.560059

**Authors:** Danyang Wang, Yongkang Tang, Ang Li, Lei Sun, Ralph E. Kleiner

## Abstract

Reactive small-molecule probes are widely used for RNA structure probing, however current approaches largely measure average RNA transcript dynamics and do not resolve structural differences that occur during folding or transcript maturation. Here, we present SNIPER-seq, an RNA structure probing method relying upon metabolic labeling with 2’-aminodeoxycytidine, structure-dependent 2’-amino reaction with an aromatic isothiocyanate, and high-throughput RNA sequencing. Our method maps cellular RNA structure transcriptome-wide with temporal resolution enabling determination of transcript age-dependent RNA structural dynamics. We benchmark our approach against known RNA structures and investigate the dynamics of human 5S rRNA during ribosome biogenesis, revealing specific structural changes in 5S rRNA loops that occur over the course of several hours. Taken together, our work sheds light on the maturation and coordinated conformational changes that take place during ribosome biogenesis and provides a general strategy for surveying evolving RNA structural dynamics across the transcriptome.

## INTRODUCTION

The folding of RNA into complex three-dimensional structures is essential for its biological function. The development of chemical methods to probe RNA structure *in vitro* and in cells has provided key insights into RNA biology. Currently, a number of different chemistries are employed to analyze RNA structure including nucleobase alkylation by DMS^1^, carbodiimides^2^ or glyoxal^3^ and 2’-hydroxyl acylation by isatoic anhydrides^4-6^ or acylimidazoles^7-9^. When combined with high-throughput RNA sequencing workflows that can detect lesion-induced misincorporation or termination during reverse transcription, chemical structure probing methods can reveal transcriptome-wide RNA structure in living cells^5, 10^. Innovations in chemistry, sequencing methods, library preparation, and bioinformatic analysis have contributed greatly to the widespread application of these methods to illuminate diverse phenomena in RNA biology^11^.

RNA structure can be modulated through binding of proteins or small molecules, alterations in environmental conditions such as temperature, oxidative stress, or solute concentration, and post-transcriptional modifications^12^. Coding and non-coding RNAs are often engaged in ribonucleoprotein (RNP) complexes, and the organization of these macromolecular assemblies, which is coordinated with RNA folding and post-transcriptional RNA processing/regulation can be highly dynamic^13^. Further, RNA folds co-transcriptionally, and therefore conditions that affect transcription can impact structural ensembles^14^. Current chemical probing methods measure the population average and obscure differences in behavior between mature RNA and newly synthesized RNA transcripts. Whereas RNA secondary and tertiary structure forms rapidly on the μs to sec timescale^15^, and is better suited for *in vitro* analyses^16^, RNA/RNP processing, maturation, trafficking, and ultimately metabolism occur at slower timescales. For example, rough estimates of ribosome biogenesis in yeast and human cells derived from pulse labeling experiments range from minutes to hours^17-19^, making this important process amenable to characterization in cellular systems. Notably, ribosome assembly pathways and intermediates in eukaryotic systems have been extensively studied using *in vitro* biochemical and structural approaches^20^, but less is known about the structure and dynamics of ribosomal biogenesis intermediates in cells. Therefore, methods to increase the temporal resolution of structure probing analyses can provide new insights into molecular mechanisms underlying the dynamic behavior and regulation of RNA in its native context.

Metabolic labeling with modified nucleosides is a powerful method to introduce a temporal dimension into studies of RNA expression and dynamics^21^. A variety of ribonucleoside analogues have been introduced to enable analysis of nascent RNA through affinity isolation or direct detection, but few have been employed in RNA structure probing workflows. Isolation of nascent RNA can be coupled with reported chemical probing strategies^22^, but this approach has not been widely used. 4-thiouridine (4-SU) and 4-thiouracil (4-TU) labeling and nascent RNA isolation have been used to facilitate DMS or hydroxyl radical probing of pre-rRNA intermediates in *E. coli* and yeast^23, 24^, but this strategy has not been applied transcriptome-wide or in human cells, and 4-SU/TU is known to perturb nucleolar morphology and rRNA processing^25^. Alternatively, the metabolic label could serve as both an affinity isolation tag and as a unique chemical handle for reactivity-based structure probing, but to our knowledge this approach has not been realized.

Here, we present synchronized nucleoside incorporation and probing evaluated by reverse transcription and sequencing (SNIPER-seq), an RNA structure probing method based upon metabolic RNA labeling with 2’-aminodeoxynucleosides combined with selective 2’-amino functionalization chemistry, enrichment, and primer extension or sequencing analysis. We demonstrate that SNIPER-seq is compatible with *in vitro* and in-cell RNA structure probing and can be applied for transcriptome-wide experiments as well as targeted analysis of non-coding RNAs. Further, SNIPER-seq enables the interrogation of RNA structure at different stages in the lifetime of a transcript, and we leverage this ability to investigate the structure of 5S rRNA during ribosomal assembly. Taken together, our strategy provides a new approach to measure transcriptome-wide RNA structure in living cells across the temporal dimension.

## RESULTS

### Quantitative analysis of 2’-amine acylation in synthetic oligonucleotides

To develop our method, we needed to identify a selective RNA functionalization reaction that reports on native structure and involves functional groups that can be introduced through metabolic labeling. Previously, Weeks and co-workers demonstrated that 2’-amino-modified RNA adopts native-like C3’-endo geometry and displays structure-dependent reactivity with NHS-esters analogous to labeling of native 2’-OH RNA with acylating agents^26^. In addition, our lab and others have previously shown that 2’-azidodeoxynucleosides can be incorporated into RNA metabolically^27, 28^ and that 2’-azido substitutions do not dramatically perturb RNA structure^29^, suggesting that small modifications at the 2’-position are metabolically tolerated and native-like. Therefore, we chose to pursue 2’-aminodeoxynucleosides as metabolically incorporated structure probes.

First, we synthesized stem-loop RNA oligonucleotides containing a single 2’-aminodeoxycytidine modification (2’-NH_2_-C) in either the stem (oligo **2**) or the loop (oligo **3**) to evaluate structure-dependent reactivity with different amine-reactive probes (Fig. 1A and Supplementary Fig. 1). A control oligonucleotide (oligo **1**) that contained only unmodified RNA was also generated. We designed three probes containing an azide reporter group for enrichment and labeling via strain-promoted azide-alkyne cycloaddition (SPAAC)^30^ and different amine-reactive warheads: 4-sulfotetrafluorophenyl ester (STP-N3), N-hydroxysuccinimidyl ester (NHS-N_3_) or aromatic isothiocyanate (isoS-N_3_) (Fig. 1B). STP, NHS, and isothiocyanates are widely used for labeling primary amines in polypeptides^31^, and sulfo-NHS^32^ and isothiocyanates^33^ have been shown to react with 2’-amine-containing RNA *in vitro* with minimal 2’-OH reactivity. Oligonucleotides (1 μM) were treated with an excess (5-20 mM) of the respective electrophilic probe for 15 min and then reacted with DBCO-biotin to generate biotinylated and mobility-shifted products (Fig. 1C). We quantified labeling efficiency of each electrophilic probe against oligos **1, 2**, and **3** by densitometry. For all three electrophilic probes, we observed a similar trend in reactivity with unpaired 2’-NH_2_-C (oligo **3**) consistently showing higher reactivity than base paired 2’-NH_2_-C (oligo **2**), and the unmodified RNA (oligo **1**) showing lowest reactivity (Fig. 1C and 1D). While labeling of unmodified oligo **1** was low using NHS-N_3_ and STP-N_3_, with 8% or 4% conversion after 20 min, respectively, labeling with isoS-N_3_ was undetectable, indicating that isothiocyanate has better specificity for the 2’-amino group (Fig. 1D). The isoS-N_3_ probe also demonstrated the highest labeling efficiency (80% conversion of oligo **3**) and comparable selectivity for unpaired 2’-NH_2_-C over base-paired 2’-NH_2_-C (Fig. 1E). We performed a time-course labeling of the three oligos with isoS-N_3_ and found that oligo **3** achieved 40% conversion in 2 min and reached a plateau after ∼15 min (Fig. 1F, Supplementary Fig. 2). Taken together, we find that the isoS-N_3_ probe displays superior reactivity and specificity for unpaired 2’-NH_2_-C nucleotides of the three probes that we assayed, and therefore it was selected for further studies.

**Figure 1.**
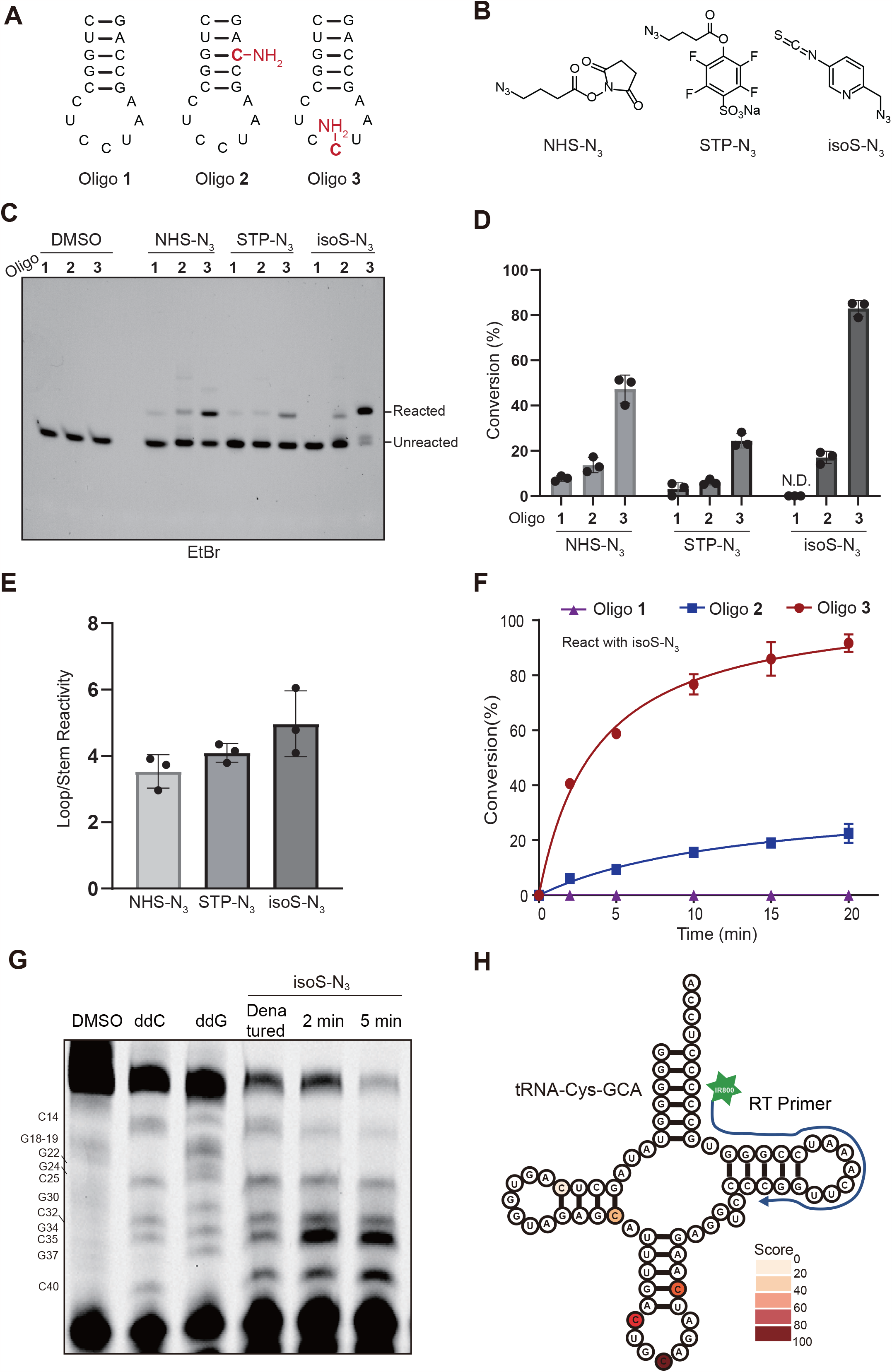
Structure-specific labeling of 2’-aminodeoxycytidine (2’-NH_2_-dCyd) in RNA oligonucleotides. **(A)** Sequence and secondary structure of oligonucleotides used in this work. 2’-NH_2_-dCyd modification is shown in red. **(B)** Chemical structure of azide-containing electrophiles used for 2’-NH_2_-dCyd labeling. **(C)** Reaction of oligo 1, 2, or 3 with azide-containing electrophiles. 1 μM oligo was reacted with 5 mM isoS-N_3_, 20 mM NHS-N_3_, or 20 mM STP-N_3_ for 15 min and then reacted with an excess of DBCO-Biotin. Reactions were separated by denaturing polyacrylamide gel and stained with EtBr. **(D)** Gel densitometry-based quantification of labeling reactions in **(C). (E)** Selectivity of each electrophilic probe for labeling structured (oligo **2**) as compared to unstructured (oligo **3**) 2’-NH_2_-dCyd sites. **(F)** Time course oligo labeling with isoS-N_3_. Reactions were performed as in **(C)**. Data represent mean ± s.d. (n = 3). **(G)** RT-primer extension-based analysis of 2’-NH_2_-dCyd-labeled IVT tRNA^Cys^ after isoS-N_3_ reaction. Nucleotide ladders were generated using ddCTP and ddGTP in the primer extension reaction. **(H)** Gel densitometry-based quantification of reverse transcription (RT) “stops” in **(G)**. RT stop scores were normalized to the intensity of the band at C35, which exhibited the most prominent stop.

**Figure 2.**
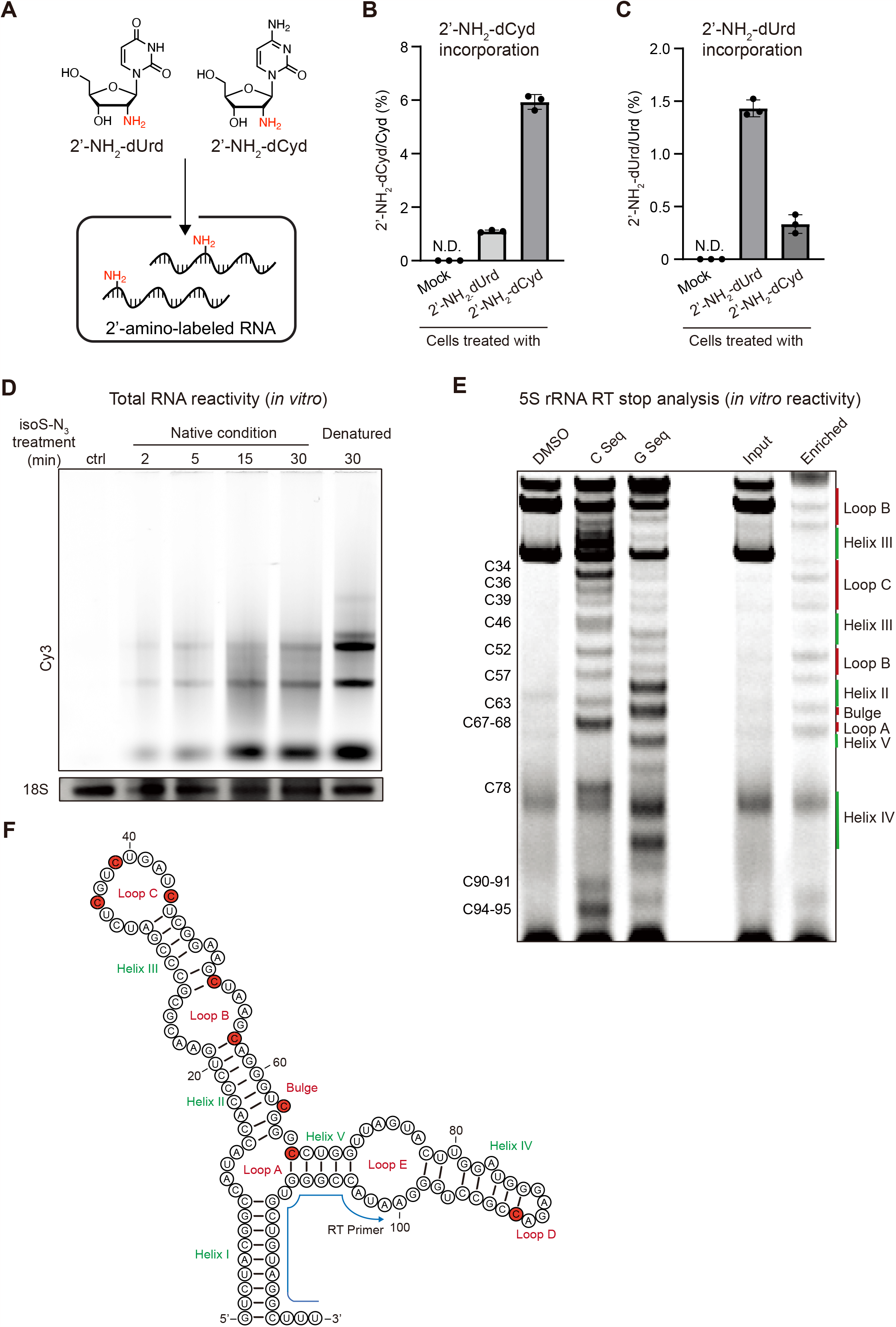
Metabolic RNA labeling with 2’-aminodeoxypyrimidines and isoS-N_3_ structure probing. **(A)**Chemical structures of nucleosides used in this work. **(B)** and **(C)** LC-QQQ-MS analysis of total RNA isolated from HEK293T treated with 1 mM 2’-aminodeoxypyrimidine nucleosides. Data represent mean ± s.d. (n = 3); n.d. = not detected. **(D)** Labeling of total RNA extracted from 2’-NH_2_-dCyd-treated HEK293T cells. Isolated total RNA was reacted with isoS-N_3_ and then treated with an excess of Cy3-DBCO. Cy3-labeled RNA was detected by in-gel fluorescence. **(E)** *In vitro* structure probing of 5S rRNA. Total 2’-NH_2_-dCyd labeled cellular RNA was purified from HEK293T cells and reacted with 5 mM isoS-N_3_ in folding buffer at 37 °C for 15 min, and then an excess of DBCO-Disulfide-Biotin for enrichment. Enriched isoS-N_3_-labeled RNA was used as template for 5S rRNA primer extension. Nucleotide ladders were generated by adding ddCTP and ddGTP to the primer extension reactions. **(F)** Secondary structure of human 5S rRNA. Residues that react with isoS-N_3_ in **(E)** are shown in red.

### 2’-NH_2_-C-based structure probing of tRNA^Cys^

We further investigated whether reaction of 2’-NH_2_-C with isoS-N_3_ could be used to measure nucleotide flexibility in a biologically relevant RNA. Based on precedent^32^, we incorporated 2’-NH_2_-C modifications into tRNA^Cys^ by *in vitro* transcription (IVT) with the corresponding modified NTP to yield, on average, a single 2’-NH_2_-C per tRNA transcript, and then measured reactivity with isoS-N_3_. We observed labeling of 2’-NH_2_-C-modified tRNA, but not unmodified tRNA, under denaturing conditions and in tRNA folding buffer (Supplementary Fig. 3), indicating that 2’-NH_2_-C is required for reactivity. Further, labeling with isoS-N_3_ was more efficient under denaturing conditions suggesting that RNA structure inhibits reaction kinetics. We next analyzed isoS-N_3_ labeling of 2’-NH_2_-C-modified tRNA^Cys^ by reverse transcription (RT)-primer extension (Fig. 1G). Consistent with known tRNA structure, we primarily observed RT termination at C residues in the anti-codon stem loop (ASL) (Fig. 1H), particularly at C35 in the anti-codon, indicating this position has the highest flexibility; in contrast, cytidine residues in the D-arm or T ΨC-arm, which are involved in tertiary interactions, displayed lower reactivity.

**Figure 3.**
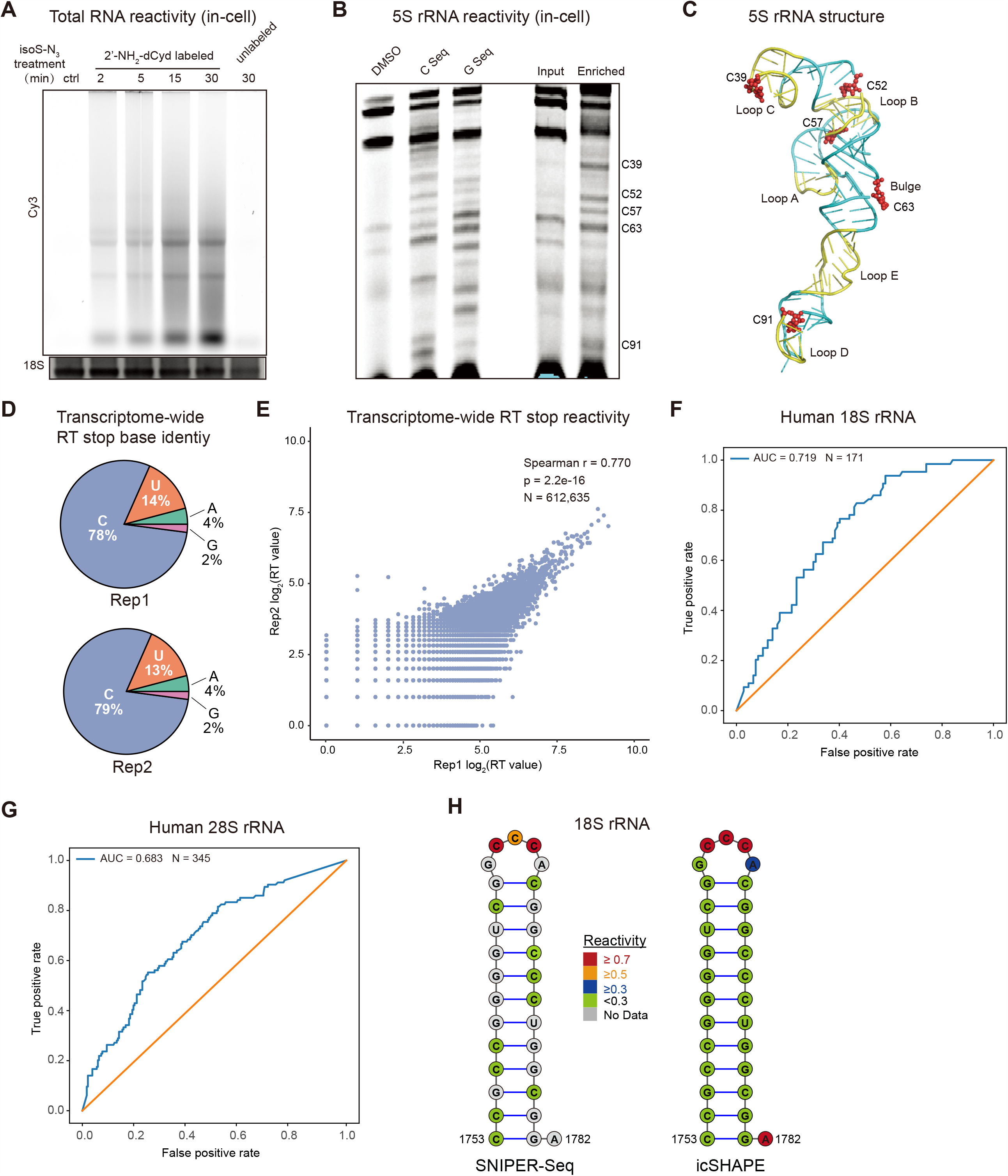
In-cell RNA structure probing with isoS-N_3_ and 2’-NH_2_-dCyd. **(A)** *In cellulo* RNA labeling with isoS-N_3_. 2’-NH_2_-dCyd treated HEK293T cells were labeled with 5 mM isoS-N_3_ in PBS at 37 °C for the indicated time. Total cellular RNA was extracted and reacted with Cy3-DBCO for detection. **(B)** *In cellulo* structure probing of 5S rRNA with isoS-N_3_. 2’-NH_2_-dCyd-labeled cells were treated with isoS-N_3_ at 37 °C for 15 min prior to RNA isolation, enrichment, and RT-primer extension analysis with a 5S rRNA-specific primer. **(C)** Structure of human 5S rRNA (PDB ID: 4UG0). Loop regions are shown in yellow and reacted cytidine residues in **(B)** are highlighted in red. **(D)** Distribution of RT stop events in two independent SNIPER-seq samples. **(E)** Scatter plot of normalized RT stop counts across the transcriptome for two independent SNIPER-seq samples; p is Spearman correlation coefficient; total number of bases = 612,635. **(F)** and **(G)** Receiver Operating Characteristic (ROC) curve plots for SNIPER-seq reactivity scores as compared against reference 18S rRNA structure (**F**) and 28S rRNA structure (**G**) from the Comparative RNA Web (CRW) site. AUC scores for all ROC curves are included. The analysis includes C residues with solvent accessibility greater than 3. **(H)** Human 18S rRNA (nucleotides 1753–1782) secondary structure with both SNIPER-seq and icSHAPE reactivity data shown in color.

### Metabolic labeling with 2’-aminodeoxynucleosides and in vitro RNA structure probing

We next applied our approach to perform structure probing on native RNA transcripts. For this purpose, we tested metabolic incorporation of 2’-aminodeoxypyrimidine nucleosides. We treated HEK293T cells with 2’-aminodeoxycytidine (2’-NH_2_-dCyd) or 2’-aminodeoxyuridine (2’-NH_2_-dUrd) for 16 hr and analyzed incorporation into total RNA by nucleoside LC-QQQ-MS (Fig. 2A, 2B, 2C, Supplementary Table 2 and 3). Gratifyingly, 2’-NH_2_-dCyd feeding resulted in 2’-NH_2_-C accumulation to 6% of unmodified C in total RNA, which is comparable or better to widely used nucleoside analogues such as 5-EU^34^ or 4-SU^35^. In addition, we also detected incorporation of 2’-NH_2_-U, likely resulting from deamination of 2’-NH_2_-dCyd. Direct feeding of 2’-NH_2_-dUrd resulted in ∼1.5% incorporation of 2’-NH_2_-U and some metabolic interconversion to 2’-NH_2_-C (Fig. 2B and 2C). Notably, neither modified nucleoside showed overt cellular toxicity (Supplementary Fig. 5) or effects on levels of native RNA pyrimidine post-transcriptional modifications (Supplementary Fig. 6). We also characterized nucleolar integrity in 2’-NH_2_-dCyd treated cells by immunostaining with NPM1 (Supplementary Fig. 7). In contrast to 4-SU, a widely used metabolic label that inhibits rRNA synthesis and induces nucleolar stress (visible by diffuse NPM1 staining), 2’-NH_2_-dCyd-treated cells showed largely unperturbed nucleolar morphology at the concentrations and treatment times evaluated.

After isolation of 2’-NH_2_-dCyd labeled total RNA, we evaluated *in vitro* labeling with isoS-N_3_, STP-N_3_, and NHS-N_3_ under denaturing conditions (Supplementary Fig. 8). While reactivity of 2’-NH_2_-modified RNA with all three probes was comparable, we observed higher background labeling of native RNA with either NHS-N_3_ or STP-N_3_, similar to our findings with synthetic oligos (Fig. 1C). We next performed a time-course analysis of the reactivity of refolded 2’-NH_2_-C-labeled total RNA with isoS-N_3_. As expected, we found that reactivity was less than under denaturing conditions but increased in a time-dependent manner from 2 min to 30 min (Fig. 2D). To verify the ability of our method to probe native RNA transcript structure, we investigated the 5S rRNA, which is highly abundant and can fold into a stable structure without the need for protein cofactors^36^. After isoS-N_3_ labeling of purified total RNA, we enriched reacted RNA through SPAAC with DBCO-disulfide-biotin and streptavidin pulldown, and measured labeling by RT-primer extension. We observed multiple RT stops corresponding to unpaired residues in 5S rRNA including bulges (C63) and loops (C26 in loop B, C34 and C39 in loop C) (Fig. 2E and 2F). Interestingly, we also found that cytidine residues at the joint of loop and stem structures showed high reactivity (i.e. C52, C57 and C67). This phenomenon was also reported when the SHAPE approach was applied to probe 5S rRNA structure^7^ and to probe structure within an RNA oligo library^37^.

### RNA structure probing in living cells using 2’-NH_2_-dCyd and isoS-N_3_

Encouraged by our *in vitro* structure probing results, we investigated whether isoS-N_3_ could probe RNA structure in living cells labeled with 2’-NH_2_-dCyd. We incubated HEK293T cells with 2’-NH_2_-dCyd overnight and treated cells with 5 mM isoS-N_3_. We then isolated total RNA and measured labeling efficiency using DBCO-Cy3. We were able to detect labeling of 2’-NH_2_-containing RNA with isoS-N_3_ as early as 2 min and reaching a plateau at ∼15 min (Fig. 3A), indicating that isoS-N_3_ is cell permeable and able to react with 2’-NH_2_-modified RNA in living cells. The labeling kinetics for isoS-N_3_ are comparable to widely used SHAPE reagents such as NAI or NAI-N_3_, albeit at 20-fold lower reaction concentration^7, 11^. We again used 5S rRNA to investigate isoS-N_3_ reaction in living cells by RT-primer extension as described above and found major RT stops at several C residues including C63, C57, C52, C39 and C91 (Fig. 3B). Analysis of the x-ray crystal structure of the human 80S ribosome, which contains the 5S rRNA and its interacting proteins (PDB ID: 4UG0), showed that these cytidines are located at loop positions and highly exposed to solvent (Fig. 3C).

We next combined metabolic labeling and isoS-N_3_ reaction with Illumina sequencing to study transcriptome-wide RNA structure (SNIPER-seq). Reaction of 2’-NH_2_-modified RNA with isoS-N_3_ was either performed *in vitro* or *in cellulo*, and unreacted native RNA was used as control. Illlumina sequencing libraries were prepared following the icSHAPE method^38^ with biotin-based enrichment of labeled RNA fragments, and isoS-N_3_ reactivity was inferred by analyzing RT stop signatures. Two independent biological replicates were used for all conditions. Compared with untreated RNA samples, in which RT stops were equally distributed across all four nucleobases, we found that 2’-NH_2_-dCyd-labeled and isoS-N_3_ treated samples predominantly contained RT stops at C (70-80% of RT stops) (Fig. 3D, Supplementary Fig. 9A). We also found that RT stops in the 2’-NH_2_-dCyd samples occurred more frequently at U (13-14%) than at A/G (6%), likely due to metabolic conversion of 2’-NH_2_-dCyd to 2’-NH_2_-dUrd, as shown previously by LC-MS analysis (Fig. 2C). Overall, >90% of RT stops occurred at C/U, demonstrating the selectivity of the isoS-N_3_ labeling reaction. Further, we observed a high correlation of RT-stop levels at individual sites between the two *in cellulo* replicates, with Spearman correlation coefficient r = 0.77, indicating that our method is robust and reproducible (Fig. 3E).

To investigate the ability our method to recapitulate complex cellular RNA structure, we chose abundant RNA transcripts with known 3D structures for analysis. We compared the reactivity profile of SNIPER-seq on all C residues in the 18S rRNA (length of 1869 nt) and 28S rRNA (length of 5348 nt) with the known structure model collected from the RiboVision database^39^, which determines secondary structure from 3D structures. SNIPER-seq reactivity profiles fit the 18S or 28S rRNA models with high area under curve (AUC) of 0.719 and 0.683, respectively (Fig. 3F and 3G). These AUC values are comparable with those obtained using icSHAPE (AUC = 0.697 for 18S rRNA, AUC = 0.633 for 28S rRNA) (Fig. 3H)^40^. In addition to rRNAs, a region of the long noncoding RNA MALAT1 was also used as a model to exam the accuracy of SNIPER-seq as its structure has been previously studied with SHAPE^41^ and it was highly abundant in our sequencing data; in this context, we also found that SNIPER-seq reactivity scores were similar to those previously determined using SHAPE and consistent with predicted structural motifs (Supplementary Fig. 9B). Taken together, our data demonstrates that SNIPER-seq can be used for *in cellulo* RNA structure probing at pyrimidine nucleosides across the transcriptome.

### Measuring transcript age-dependent structural changes with SNIPER-seq

We explored whether we could use SNIPER-seq to study time-dependent structural changes occurring in RNA in a pulse-chase experiment with 2’-NH_2_-dCyd (Fig. 4A). The biogenesis of ribosomes involves the synthesis, processing, and assembly of multiple RNA transcripts including 28S, 18S, 5.8S, and 5S rRNA^20^. 28S, 18S, and 5.8S rRNAs are processed from a single 45S rRNA transcript generated in the nucleolus by RNA Pol I, whereas the 5S rRNA is transcribed from a separate locus (outside of nucleoli) by RNA Pol III before it forms the 5S ribonucleoprotein particle (5S RNP) and is imported into the nucleolus and incorporated into the pre-60S particle. The 5S RNP then undergoes 180 degree rotation to form the central protuberance (CP) structure that is found in the mature 60S subunit. Whereas the order of these steps in yeast and human ribosome biogenesis have been established through *in vitro* structural studies^42, 43^, their timing in cells and the native distribution of intermediates is poorly established. We fed cells 2’-NH_2_-dCyd for 1 hr to label newly transcribed RNA, performed a chase for 3, 6, or 12 hr, and then reacted cells with isoS-N_3_ at the end of the chase. We first analyzed isoS-N_3_ labeling by performing targeted RT-primer extension of 5S rRNA. Interestingly, we observed differences in reactivity at different chase time points (Fig. 4A). In particular, reactivity at positions C39 in loop C and C91 in loop D increased as a function of chase time, while labeling at C34 in loop C and C78 in helix V decreased in a time-dependent manner. In contrast, the reactivity at positions C52, C57, and C63 remained constant over the course of our experiment.

**Figure 4.**
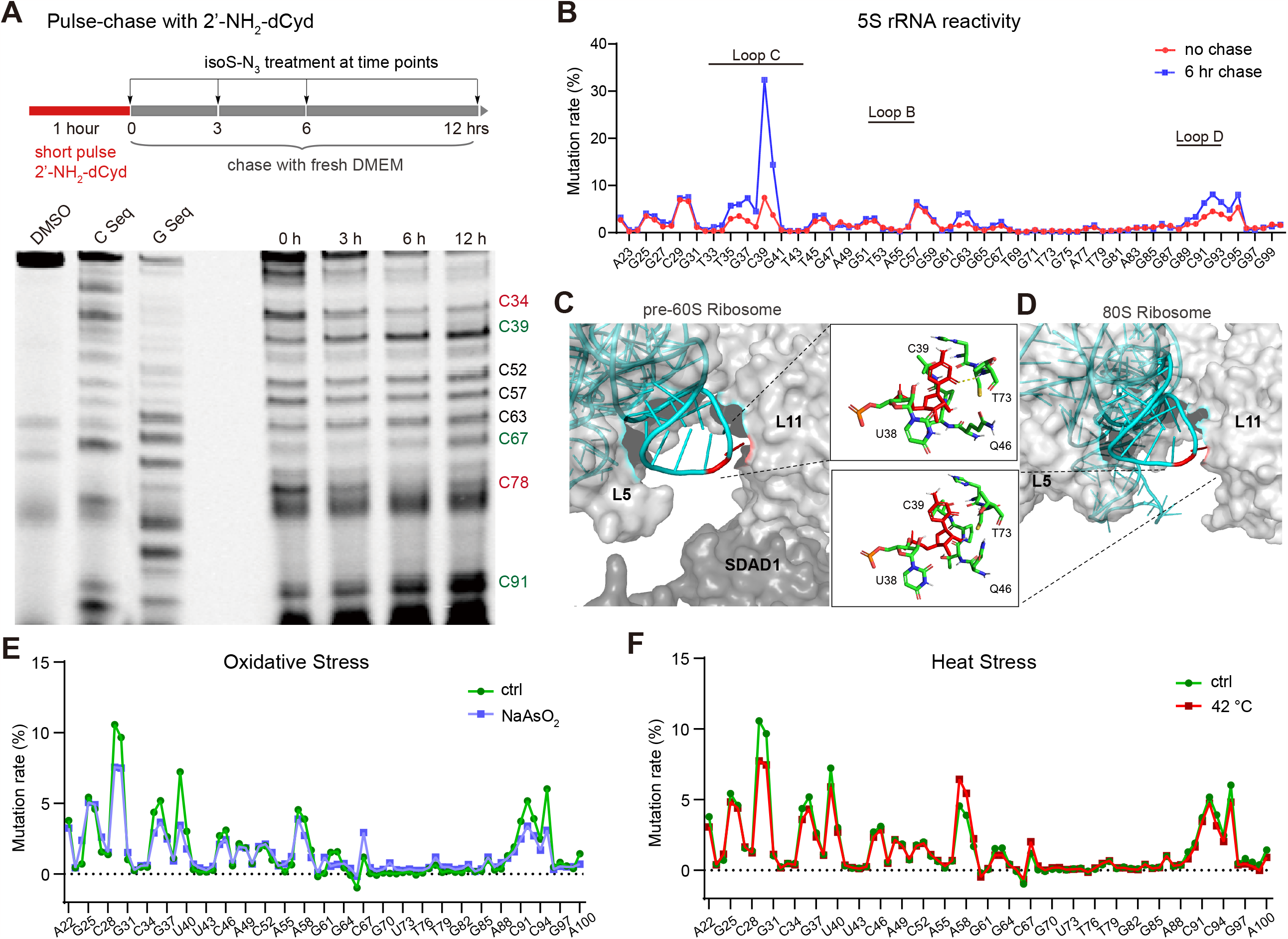
In-cell nascent RNA structure probing with SNIPER-seq. **(A)** Probing time-dependent 5S rRNA structural dynamics using pulse-chase feeding with 2’-NH_2_-dCyd and isoS-N_3_ labeling. Enriched isoS-N_3_ labeled RNA was used for 5S rRNA primer extension reaction. **(B)** Mutational profiling to measure 5S rRNA structural dynamics. Cells were pulsed for 1 hr with 2’-NH_2_-dCyd and processed immediately or chased for 6 hr. **(C)** and **(D)** Structure of 5S rRNA loop C (cyan) in complex with ribosomal proteins (gray) from human pre-60S ribosome (PDB ID: 3JCT) **(C)** or mature 80S ribosome (PDB ID:4V7R) **(D)**. C39 residue is shown in red and interactions with proximal protein residues are shown in the inset. **(E)** and **(F)** Structural dynamics of nascent 5S rRNA during oxidative stress **(E)** or heat stress **(F)**. For oxidative stress, cells were cotreated with 0.5 mM NaAsO_2_ and 1 mM 2’-NH_2_-dCyd for 1 hr at 37 °C. For heat stress, cells were incubated with 1 mM 2’-NH_2_-dCyd at 42 °C for 1 hr.

To further validate differences in nucleotide reactivity on the 5S rRNA, we measured adduct formation by mutational profiling immediately after 1 hr 2’-NH_2_-dCyd pulse labeling or after 1 hr pulse labeling followed by a 6 hr chase. We reverse transcribed RNA in the presence of Mn^2+^, a condition known to induce read through with mutation at modified nucleotide adducts^44, 45^, and performed high-throughput sequencing of the 5S rRNA amplicon. Total mutation rate at each nucleotide was calculated by combining the frequency of substitutions, deletions, and insertions and subtracting the mutation rate present in the control. In the 1 hr pulse sample, we found mutation rates above 5% at residues C29, C30, C39, C57, C91 and C95 (Fig. 4B and Supplementary Fig. 10). After a 6 hr chase (Fig. 4A), the most dramatic change in reactivity occurred at C39 in loop C, which showed a 4-fold increase in mutational frequency (this position also showed a time-dependent increase in reactivity in the gel-based RT stop analysis). We also found a time-dependent 1.7-fold increase in mutation rate at C36 in loop C and a 1.6-fold increase in mutation at C95 in loop D. In contrast, mutational frequencies at C30 in helix III and C57 in loop B remained largely unchanged during the chase, suggesting that they undergo only minimal changes in structure and reactivity. To understand the origin of reactivity differences at C39 in loop C, we examined structures of the 5S RNP from pre-60S subunit intermediates to the mature 80S ribosome in yeast and human^42, 43^. Whereas the secondary and tertiary structure of the 5S rRNA is largely conserved throughout ribosome biogenesis, (Supplementary Fig. 11A), its interaction with ribosomal proteins is dynamic. During the initial association of 5S RNP with the pre-60S ribosome (State I1, PDB ID: 8FL2)^42^, loop C is situated within a binding pocket formed by two ribosomal protein, L5 and L11, and nucleolar protein SDAD1 (Fig. 4C). After 180 degree rotation and in the mature 60S/80S (PDB ID: 4UG0)^46^, loop C becomes more solvent exposed (Fig. 4D). We also found a hydrogen bond between the O2 atom of C39 and amino acid residue Thr73 in L11 within the pre-rotated state (Fig. 4C, inset). After 5S RNP rotation, C39 drifts apart from L11, losing the hydrogen bond between C39 and Thr73 (Fig. 4D, inset). We propose that the increased solvent accessibility and loss of hydrogen bond in the mature 5S RNP state are responsible for the increased reactivity in loop C observed in our SNIPER-seq experiment. Notably, we also found increased solvent accessibility in this region in structural intermediates from yeast ribosomal assembly (Supplementary Fig. 11B, 11C)^43, 47^.

We were also curious how the folding and maturation of nascent rRNA is affected by stress conditions. Oxidative stress can affect rRNA processing^48^ and heat stress has been shown to unfold mRNA^49^ and tRNA^50^ structure. We performed SNIPER-seq on newly transcribed 5S rRNA under oxidative stress (NaAsO_2_) or heat stress conditions together with mutational profiling (Fig. 4E, 4F). Compared to the unstressed control, oxidative stress reduced isoS-N_3_ reactivity at multiple sites, including C28 in helix III, C36 and C39 in loop C, and C91 in loop D (Fig. 4E, Fig. 2F). The reactivity of C26 and C57 in loop B remained at the same level. In contrast, only C28 showed decreased reactivity under heat stress, whereas C57 exhibited slightly higher reactivity. Upon nucleolar stress, such as heat shock, serum starvation and oxidative stress, integration of 5S RNP into the pre-60S is decreased^51^, and 5S RNP accumulates in the nucleoplasm, forming interactions with the E3 ubiquitin ligase HDM2^52^. These processes may collectively contribute to the observed differences in 5S rRNA structural probing results under stress conditions. Taken together, our study demonstrates how age-dependent changes in cellular RNA structure can be revealed by SNIPER-seq, providing a versatile method to study RNP maturation and condition-dependent structural changes.

## DISCUSSION

In this study, we present a novel method for probing RNA structure in living cells with the ability to selectively study RNA transcripts as a function of their lifetime. Our method, SNIPER-seq, relies upon metabolic labeling of cellular RNA with 2’-NH_2_-dCyd, which is incorporated at high levels through native metabolic pathways with minimal cytotoxicity. Metabolic labeling facilitates temporal resolution of specific RNA populations and the 2’-amino modification serves as a non-perturbing structure-dependent reactivity handle for labeling and enrichment with a cell-permeable isothiocyanate probe. We validate our method using RNAs of known structure and analyze the age-dependent structure of 5S rRNA over the course of ∼1-12 hr after its transcription to reveal dramatic and region-specific structure and reactivity differences in this RNA species during its lifetime.

SNIPER-seq endows reactivity-based RNA structure probing methods with temporal resolution. In this work we demonstrate how our method can be used to study the dynamics of RNA maturation within the context of ribosome biogenesis. Moving forward, we envision broad application of this approach to study age-dependent RNA dynamics transcriptome-wide on coding and non-coding RNAs. Whereas our method may currently lack the ability to characterize co-transcriptional folding (which typically occurs on the μs-sec time scale), structural dynamics that result from changes in protein association, complex assembly, or cellular localization during the lifetime of an RNA should be amenable to our approach, thereby establishing the absolute timing of these processes. Further, pulse-labeling allows the specific interrogation of perturbations to co-transcriptional folding or RNA processing pathways that result in longer-lived structural intermediates that can be probed. Our method therefore provides a complementary approach to structure probing methods that operate on the entire transcriptome and should provide new insights into RNA structure and interactomes in cells.

We applied SNIPER-seq to study the age-dependent dynamics in the 5S rRNA. We measure changes in RNA reactivity at specific loop residues that occur over the span of multiple hours, such as the dramatic increase in reactivity in loop C. RNA pulse labeling in human cells has established rough estimates for rRNA maturation^17, 19^, however the absolute timing of specific events and intermediates in human ribosome biogenesis are still largely unknown. Our study suggests that conformational changes in the 5S rRNA occur later than estimates for maturation of the 18S and 28S rRNA. In contrast to other canonical rRNAs that are generated from the 45S rRNA, 5S is transcribed from a separate locus outside of the nucleolus by RNA Pol III and incorporated into the pre-60S subunit later as the 5S RNP, consistent with slower maturation kinetics. Alternatively, the observed structural dynamics for 5S may reflect conformational changes occurring in cytosolic ribosomes.

Metabolic labeling with modified nucleosides has been widely used to install orthogonal chemical functionalities for RNA detection and affinity isolation. Here we use this approach to directly incorporate an artificial functional group that reports on RNA structure. Whereas the 2’-amino group is metabolically tolerated and compatible with native RNA structure, its reactivity is greatly enhanced relative to 2’-OH or any of the nucleobase exocyclic amines present on RNA, which enables selective functionalization of modified nucleotides. Here we present our strategy using 2’-NH_2_-dCyd, but it is likely that other 2’-amino modified nucleosides will also be compatible with our approach, expanding its utility to probe all 4 nucleotides; 2’-thionucleosides have also been incorporated into RNA transcripts^53^ and are likely compatible with cellular metabolism and our approach. Further, we envision that perturbation of nucleotide metabolism to increase flux through the nucleoside salvage pathways may offer a strategy to enhance nucleoside uptake and provide even higher temporal resolution. Such studies are underway in our group and will be reported in due course.

## ACKNOWLEDGMENTS

We thank D. Dhinghani for assistance with bioinformatic analysis. R.E.K. acknowledges support from the NIH (R01 GM132189) and NSF (MCB-1942565). R.E.K., D.W., and A.L. acknowledge financial support from Princeton University.

