## Supplementary Information for "Transcriptome-wide RNA structure probing with temporal resolution"

Contents

Materials and Methods 2

Supplementary Tables 12

Supplementary Figures 15

Synthetic Methods and Compound Characterization 26

### Materials and methods

*Chemicals and Synthesis*

Starting materials and reagents were purchased at ACS reagent grade or higher from Acros Organic or Sigma-Aldrich and used without further purification. Analytical thin-layer chromatography was performed using pre-coated MilliporeSigmaTM 200 µm silica gel F-254 plates. Bands were visualized by UV and stained with Hanessian's stain (cerium molybdate). $^{1}$H NMR spectra were recorded on a 500 MHz Bruker spectrometer using an AutoX PFG probe at 298 K; residual solvent peaks were used as internal references: DMSO (quint, δH = 2.50 ppm), $CHCl_3$ (s, δH = 7.26 ppm) or methanol (quint, δH = 3.31 ppm). $^{13}$C NMR spectra were recorded on a 500 MHz Bruker spectrometer using an AutoX PFG probe at 298 K; residual solvent peaks were used as internal references: DMSO (δ = 40.50 ppm), $CHCl_3$ (δ = 77.23 ppm) or methanol (δ = 49.00 ppm). Coupling constants (J) are reported in hertz (Hz). The following abbreviations are used to describe the multiplicities: s = singlet, d = doublet, t = triplet, q = quartet, quint = quintet, sext = sextet, m = multiplet, dd = doublet-doublet.

*Oligonucleotide synthesis*

Oligonucleotide synthesis was performed on an Applied Biosystems (ABI) 394 oligonucleotide synthesizer using standard coupling conditions and commercial oligo synthesis reagents and phosphoramidites (Glen Research) unless otherwise noted. 2'-aminodeoxycytidine phosphoramidite was purchased from ChemGenes. Oligos were cleaved from the resin and deprotected under standard conditions. Oligos were purified by 20% denaturing urea-polyacrylamide gel.

*$tRNA^{Cys}$ in vitro transcription*

$tRNA^{Cys}$ DNA template was synthesized using Klenow DNA polymerase (NEB), with overlapping primers (Supplementary Table 1) and purified by 3% agarose gel. IVT $tRNA^{Cys}$ was generated with T7 RNA polymerase in IVT buffer (300 μL) containing 50 mM Tris (pH 7.5), 15 mM $MgCl_2$, 5 mM DTT, 2 mM spermidine, 10 pmol DNA template and 2 mM of each nucleotide triphosphate. The reaction was incubated at 37 °C for 2 hours. To generate tRNA containing 2'-aminodeoxycytidine substitutions, the transcription reaction contained 2 mM ATP, 2 mM GTP, 2 mM UTP, 1.2 mM 2'-aminodeoxycytidine triphosphate, and 0.8 mM CTP. RNA was purified by 8% denaturing urea-polyacrylamide gel.

*Oligo Labeling*

Synthetic RNA oligos (1 μM) were labeled with 20 mM NHS-$N_3$, 20 mM STP-$N_3$ or 5 mM isoS-$N_3$ in 20 μL folding buffer (100 mM HEPES pH 8.0, 6 mM $MgCl_2$ and 100 mM NaCl) for the indicated time. Reactions were quenched by adding 1 μL of 1 M hydroxylamine, and purified by ethanol precipitation. The labeled oligo was further reacted with 50 μM DBCO-Biotin at 37 °C for 2 hours in 1X PBS (phosphate buffered saline) solution. Products were resolved by 15% denaturing polyacrylamide gel. Band intensities were quantified using ImageJ.

$tRNA^{Cys}$ (10 pmol) was dissolved in 45 μL folding buffer and incubated at 37 °C for 15 min. Next, 5 μL isoS-$N_3$ (50 mM in DMSO) was added into the reaction and incubated for the indicated time. For labeling under denaturing conditions, $tRNA^{Cys}$ was dissolved in folding buffer/formamide (1:1) solution and then reacted with isoS-$N_3$. The reaction was quenched with 1 μL of 1 M hydroxylamine and purified by ethanol precipitation. The labeled $tRNA^{Cys}$ was used as a template for reverse transcription or further reacted with 50 μM DBCO-Cy3 for detection.

*Primer extension*

300 ng labeled tRNA$^{Cys}$ was mixed with 3 pmol IR-800 labeled RT Primer (Supplementary Table 1) in 15 μL $H_2O$ and 1 μL 10 mM dNTP mix was added. For generation of a dideoxy sequencing ladder, 1 μL 10 mM dNTP and 4 μL 10 mM ddNTP were used in the primer extension reaction. The reaction was incubated at 65 °C for 5 min and then cooled to 4 °C at a rate of 0.2 °C /s. Next, 4 μL 5X First-Strand Buffer (Invitrogen), 1 μL 0.1 M DTT, and 0.25 μL SuperScript III (Invitrogen) were added to a final volume of 20 μL. The reaction was incubated with the following conditions: 25 °C for 10 min and 52 °C for 1 hour. 20 μL loading dye was added and the mixture was denatured at 95 °C for 3 min before separation by denaturing 8% polyacrylamide gel and visualization by in-gel fluorescence scanning.

*Cell culture and metabolic labeling*

HEK293T cells were cultured at 37 °C in a humidified atmosphere with 5% $CO_2$ in DMEM (Life Technologies) supplemented with 10% fetal bovine serum (Atlanta), 1x penicillin-streptomycin (Life Technologies) and 2 mM L-glutamine (Life Technologies). For nucleoside labeling experiments with HEK293T cells, cells were seeded at $3 \times 10^6$ cells in a 10 cm petri dish. 24 hours later, fresh medium containing 1 mM 2'-$NH_2$-dUrd or 2'-$NH_2$-dCyd was added and incubated for the indicated time.

*Nucleoside LC-QQQ-MS*

HEK293T cells treated with 1 mM 2'-$NH_2$-dUrd or 2'-$NH_2$-dCyd were harvested and total RNA was extracted using Trizol reagent (Invitrogen) according to the manufacturer's instructions. RNA samples (2 μg) were digested with nuclease P1 (Wako Chemicals, 2U) in 30 μL of buffer

(7 mM NaOAc, 0.4 mM $ZnCl_2$, pH 5.2) at 37 °C for 2 hours. The mixture was dephosphorylated with Antarctic Phosphatase (NEB, 1 µL) at 37 °C for 2 hours.

UHPLC-MS/MS analysis of digested RNA samples was performed on an Agilent 6470 triple quadrupole LC/MS system. The following source parameters were used for the mass spectrometry: gas temperature 175 °C, gas flow 12 L/min, nebulizer 20 psi, sheath gas temperature 325 °C, sheath gas flow 12 L/min, capillary voltage 2,500 V in the positive ion mode, capillary voltage -2,500 V in the negative ion mode. Chromatography was performed on a Thermo Scientific Hypersil GOLD aQ column (3 µm, 150 x 2.1 mm) at 36 °C using water (containing 0.1% formic acid) at a flow rate of 0.4 mL/min. The injection volume was 1 µL. The mass transitions and retention times shown in Supplementary Table 2 were used to identify each nucleoside. Calibration curves for quantification were generated by injection of various concentrations of nucleoside standards (Supplementary Figure 4).

*Total RNA in vitro labeling*

Total cellular RNA was extracted from 2'-NH2-dCyd treated HEK293T cells. 10 µg total RNA was labeled with 20 mM NHS-$N_3$, 20 mM STP-$N_3$, or 5 mM isoS-$N_3$ at 37 °C for 15 min in 20 µL folding buffer (100 mM HEPES pH 8.0, 6 mM $MgCl_2$ and 100 mM NaCl). RNA labeling under denaturing conditions was performed in folding buffer containing 50% formamide. Reactions were quenched with 1 µL 1 M hydroxylamine. The reaction was purified with Zymo RNA clean and concentrator-5 spin columns according to the manufacturer's instructions, and then reacted with 50 µM Cy3-DBCO using SPAAC in PBS solution at 37 °C for 2 hours. After Zymo column purification, labeled total RNA was resolved by native gel electrophoresis using 1% TAE-agarose gel. Labeled RNA was visualized by in-gel fluorescence on a

TyphoonFLA 9500 Fluorescent Image Analyzer Scanner (GE Healthcare) using a Cy3 filter set. Total RNA was visualized by staining with EtBr staining as loading control.

*In-cell labeling of RNA*

HEK293T cells treated with 1 mM 2'-$NH_2$-dCyd were rinsed once with 1× PBS and scraped off the plate in 5 mL PBS. After centrifugation, the cell pellet was resuspended in 900 µL PBS. 100 µL freshly prepared isoS-$N_3$ (50 mM in DMSO) was added slowly to cells and incubated at 37 °C for the time indicated. The reaction was quenched with 10 uL 1M hydroxylamine and the cell pellet was collected by centrifuge. Total RNA was extracted and reacted with DBCO-Cy3 as described above.

*Enrichment of isoS-$N_3$ labeled RNA*

Total cellular RNA containing 2'-$NH_2$-dCyd and reacted with isoS-$N_3$ either *in vitro* or *in cellulo*, was further reacted with 100 µM DBCO-Disulfide-Biotin using SPAAC in PBS solution at 37 °C for 2 hours. After ethanol precipitation, 100 µg total RNA was incubated with 100 µL prewashed Dynabeads™ M-280 Streptavidin beads for 1 hr at room temperature. Next, beads were washed three times with 0.5 x B&W (2.5 mM Tris pH 7.5, 0.25 mM EDTA, 500 mM NaCl) buffer, and RNA was eluted by treating the beads with 100 mM DTT at 37 °C for 30 mins. The elution was further purified using Zymo RNA clean and concentrator-5 spin columns according to the manufacturer's instructions.

*5S rRNA primer extension*

2 µg enriched total RNA was mixed with 3 pmol IR-800 labeled RT Primer (Supplementary Table 1) in 15 µL and 1 µL 10 mM dNTP mix was added. For generation of a dideoxy sequencing ladder, 2 µg total RNA was used as template and 1 µL 10 mM dNTP and 4 µL 10

mM ddNTP were added into the reaction. The reaction was incubated at 65 °C for 5 min and then cooled to 4 °C at a rate of 0.2 °C /s. Next, 4 µL 5X First-Strand Buffer (Invitrogen), 1 µL 0.1 M DTT, and 0.25 µL SuperScript III (Invitrogen) were added to a final volume of 20 µL. The reaction was incubated with the following conditions: 25 °C for 10 min and 52 °C for 1 hr. 20 µL loading dye was added, the mixture was denatured at 95 °C for 3 min, and cDNA was resolved on a denaturing 8% polyacrylamide gel and visualized by in-gel fluorescence.

*RNA sequencing library preparation*

Transcriptome-wide RNA sequencing was performed with two independent biological replicates per condition. Three conditions were assayed: DMSO treated, *in vitro* isoS-$N_3$ reaction on purified RNA, and *in cellulo* isoS-$N_3$ reaction. For *in vitro* labeling, total cellular RNA extracted from 2'-$NH_2$-dCyd-treated HEK293T cells was reacted with isoS-$N_3$ in folding buffer as described above. For *in cellulo* treated samples, 2'-$NH_2$-dCyd labeled HEK293T cells (1 mM for 16 hr) were treated with 5 mM isoS-$N_3$ for 15 min as described above and RNA was isolated with Trizol reagent. Poly(A)+ RNA was isolated using oligo(dT) beads starting with 200 µg isoS-$N_3$ labeled total RNA. After isoS-$N_3$ labeling, RNA was reacted with 100 µM DBCO-Biotin for 2 hr at 37 °C in PBS. After Zymo column purification, poly(A)+ RNA was fragmented with fragmentation reagent (Thermo Fisher) at 90 °C for 1 min. Fragmented poly(A)+ RNA was end-repaired using 1 U FastAP and 10 U T4 PNK in 10 µl T4 PNK buffer at 37 °C for 1 h. After 3' end dephosphorylation, the RNA was ligated to the 3' adaptor-ddC (Supplementary Table 1) at 16 °C overnight with T4 RNA ligase I (NEB). For the DMSO control sample, RNA was ligated to 3' adaptor-biotin (Supplementary Table 1). The ligation product was purified by 8% denaturing urea-polyacrylamide gel. Bands above 50 nt were excised and purified. Purified RNA samples were incubated with 1 µL of 10 µM icSHAPE-RT primer (Supplementary Table 1) and 1 µL of 10 mM dNTPs in 20 µL reaction volume. Next, samples were heated to 70 °C for 5 min and then cooled slowly to 25 °C (0.2 °C/s) and incubated at 25 °C for 1 min. After primer annealing, 0.5 μL of SUPERaseIn, 1 μL 100 mM DTT, 4 μL of 5× First Strand Buffer (Thermo Fisher) and 1 μL of SuperScript III (Thermo Fisher) were added. Reverse transcription was performed at 25 °C for 3 min, 42 °C for 5 min, and 52 °C for 30 min. After cDNA extension, samples were kept below 37 °C to avoid denaturing RNA-cDNA hybrids. To the 20 μL reverse transcription reaction was then added 10 μL pre-washed Dynabeads™ MyOne™ Streptavidin C1 solution, and the mixture was incubated at 25 °C for 45 min with rotation. The beads were washed 4x with 500 μL of 4 M NaCl wash buffer (100 mM Tris pH 7.0, 4 M NaCl, 10 mM EDTA and 0.2% Tween 20) and cDNA was eluted with RNaseA/T1/H mix at 37 °C for 30 min. cDNA was circularized by CircLigase II (Epicentre) and then amplified using Solexa primers and submitted for Illumina sequencing.

*Bioinformatic analysis*

SNIPER-Seq analysis was performed following the icSHAPE protocol, and we employed a congruous approach for processing the sequencing reads. This involved employing the icSHAPE-pipe scripts accessible at (https://github.com/lipan6461188/icSHAPE-pipe). Furthermore, to ensure comprehensive coverage, we also made adaptations to the Demultiplexer_v2.py script (https://github.com/GrosseLab/iCLIP/blob/master/Demultiplexer_v2.py) to split raw fastq files based on the associated barcodes (Supplementary Table 1). Subsequently, we executed icSHAPE-pipe's trim function to excise any potential adapter sequences. This operation was carried out with specific parameters (-l 13 -a adaptor_zhangqc.fa), and the adapter sequence can be found here: https://github.com/qczhang/icSHAPE/tree/master/data/adapter.

Refined reads were first mapped to ribosomal RNAs as well as small RNAs (with lengths less than 200). For reads that persisted as unmapped in this initial phase, we proceeded with their alignment to the human genome (hg38) using STAR in the end-to-end mode. Following mapping, we computed RT-stop signals by employing the sam2tab utility from the icSHAPE-pipe toolkit, which was subsequently followed by utilizing the calcSHAPE module, also from icSHAPE-pipe, to derive the final RT-stop signals. We calculated SHAPE scores through the application of icSHAPE-pipe's genSHAPEToTransSHAPE function.

To perform comparative analysis between SNIPER-Seq and known RNA structures, we procured the secondary structure models for human 18S rRNA and 28S rRNA from a cryo-EM structure of the human 80S ribosome (PDB ID: 6EK0). This enabled us to calculate the solvent accessibility of each nucleotide within the 3D model, retaining those bases with a solvent accessibility greater than 3. This assessment was integral for evaluating the Area Under the Curve (AUC) metric. The reactivity score profiles for SNIPER-Seq were recalculated upon remapping to these rRNA structures.

To gauge the congruence between the structural probing reactivity scores and the reference structure model, we generated Receiver Operator Characteristic (ROC) curves. These curves quantified the extent to which the reactivity scores aligned with the anticipated structural model. By employing varying reactivity score cutoffs, we classified each nucleotide as either single-stranded or double-stranded. Here, a true positive was defined as a single-stranded base with a reactivity score surpassing the threshold, whereas a true negative referred to a paired base with a reactivity score lower than the cutoff. Importantly, only C nucleotides exhibiting an accessibility score greater than three were retained for this evaluation.

We accessed icSHAPE data from the Gene Expression Omnibus under the accession number GSE153984. To visually depict the RNA structures, we utilized the VARNA tool.

*Pulse chase and amplicon sequencing*

HEK293T cells were treated with 1 mM 2'-$NH_2$-dCyd for 1 hour, and subsequently chased for the indicated time in fresh DMEM. For the *in cellulo* isoS-$N_3$ reaction, total RNA extraction and enrichment of isoS-$N_3$ labeled RNA were performed as described above. For control samples, we omitted 2'-$NH_2$-dCyd feeding or the isoS-$N_3$ labeling step (results were comparable). 1 µg enriched RNA or control total RNA was annealed with 2 pmol 5S rRNA primer in 11 µL $H_2O$ at 65 °C for 5 min and then cooled on ice. 8 µL of 2.5× MaP buffer (125 mM Tris-HCl pH 8.0, 1.25 mM dNTPs, 188 mM KCl, 25 mM DTT, and 15 mM $MnCl_2$) was added to each tube. The reaction mix was incubated at 42 °C for 2 minutes. Then, 1 µL SuperScript II (Thermo Fisher) was added to the reaction mix. The reverse transcription mix was incubated at 42 °C for 3 h. The reaction was terminated by heating at 70 °C for 15 min, and the cDNA was purified with a DNA Clean & Concentrator-5 columns (Zymo). cDNA was amplified with Phusion HF DNA polymerase (NEB). PCR products were purified with a 2% agarose gel and submitted to Genewiz for amplicon sequencing. The sequencing data were aligned and analyzed with the CRISPResso2 tool (http://crispresso.pinellolab.org/).

*Immunofluorescence*

HEK293T or HeLa cells were seeded on 12 mm coverslips (Fisher Scientific, 12-545-81) in 24-well plates. Cells were treated with either 200 µM 4-thiouridine or 1 mM 2'-$NH_2$-dCyd for the indicated time. Following incubation, cells were washed once with PBS, fixed with PBS containing 3% paraformaldehyde at 37 °C for 20 minutes, and washed once with PBS. Next, cells were permeabilized with 0.5% Triton-X in PBS for 10 min at RT. The fixed cells were blocked with 5% goat serum in PBS for 1 hr, then incubated with NPM1 monoclonal antibody

(Invitrogen, NA-24, 1:300 dilution) for 2 hr at RT. After washing three times with PBS, the cells were incubated with Goat anti-mouse Alexa 488 antibody (Jackson Immuno Research, 1µg/mL) for 1 hr at RT in the dark. The coverslips were then washed twice more with PBS for 5 min each, stained with Hoescht33342 (Thermo Scientific,1 ug/mL) for 10 min, washed with PBS once more, and mounted on glass microscopy slides in ProLong Anti-Fade Reagent (Life Technologies) and sealed with nail polish.

**Supplementary Table 1.** Oligonucleotides used in this work

| Name | Sequence |
|---|---|
| $tRNA^{Cys}$ template-for | AAGCTTAATACGACTCACTATAGGGGGTATAGCTCAGTGGTAGAGCATTTGACTGCAGAT |
| $tRNA^{Cys}$ template-rev | AGGGGGCACCCGGATTTGAACCGGGGACCTCTTGATCTGCAGTCAAATGCTCTACCACTG |
| $tRNA^{Cys}$ RT Primer | IR800-GGCACCCGGATTTGAACCG |
| 5S rRNA RT Primer | IR800-AAAGCCTACAGCACCCGGTAT |
| 3'Adaptor-ddC | /5rApp/AGATCGGAAGAGCGGTTCAG/3ddC/ |
| 3'Adaptor-Biotin | /5rApp/AGATCGGAAGAGCGGTTCAG/3Biotin/ |
| icSHAPE RT-1 | /5phos/DDDNNGGTTNNNNAGATCGGAAGAGCGTCGTGGA/iSp18/GGATCC/iSp18/TACTGAACCGC |
| icSHAPE RT-2 | /5phos/DDDNNTTGTNNNNAGATCGGAAGAGCGTCGTGGA/iSp18/GGATCC/iSp18/TACTGAACCGC |
| icSHAPE RT-3 | /5phos/DDDNNACCTNNNNAGATCGGAAGAGCGTCGTGGA/iSp18/GGATCC/iSp18/TACTGAACCGC |
| icSHAPE RT-4 | /5phos/DDDNNCAATNNNNAGATCGGAAGAGCGTCGTGGA/iSp18/GGATCC/iSp18/TACTGAACCGC |
| icSHAPE RT-5 | /5phos/DDDNNTGGCNNNNAGATCGGAAGAGCGTCGTGGA/iSp18/GGATCC/iSp18/TACTGAACCGC |
| icSHAPE RT-6 | /5phos/DDDNNGGTCNNNNAGATCGGAAGAGCGTCGTGGA/iSp18/GGATCC/iSp18/TACTGAACCGC |
| 5S rRNA RT Primer | AAAGCCTACAGCACCCGGTAT |
| 5S rRNA PCR primer-for | ACACTCTTTCCCTACACGACGCTCTTCCGATCTGTCTACGGCCATACCACCCTG |
| 5S rRNA PCR primer-rev | GACTGGAGTTCAGACGTGTGCTCTTCCGATCTAAAGCCTACAGCACCCGGTAT |

**Supplementary Table 2**. LC-QQQ-MS parameters used for measurement of modified and canonical nucleosides in total cellular RNA. CE: collision energy, CAV: collision cell accelerator voltage. Nucleosides were detected in positive ion dynamic multiple reaction monitoring (DMRM) mode.

| **Compound** | **Precursor Ion (m/z)** | **Product Ion (m/z)** | **Fragmentor** | **CE (V)** | **CAV (V)** | **Polarity** |
|---|---|---|---|---|---|---|
| 2'-$NH_2$-dCyd | 243 | 112 | 166 | 20 | 5 | Positive |
| 2'-$NH_2$-dUrd | 244 | 113 | 166 | 14 | 5 | Positive |
| Cyd | 244 | 112 | 166 | 20 | 5 | Positive |
| Urd | 245 | 113 | 166 | 14 | 5 | Positive |
| D | 247 | 115 | 166 | 14 | 5 | Positive |
| $m^5C$ | 258 | 126 | 166 | 14 | 5 | Positive |

**Supplementary Table 3**. LC-QQQ-MS analysis of total RNA from HEK 293T cells treated with 2'-$NH_2$-dCyd/dUrd. Values are reported as ng/mL except where indicated.

| | Cyd | Urd | 2'-$NH_2$-dCyd | 2'-$NH_2$-dUrd | D | $m^5C$ | 2'-$NH_2$-dCyd/ Cyd (%) | 2'-$NH_2$-dUrd/ Urd (%) | D/Urd (%) | D/Cyd (%) |
|---|---|---|---|---|---|---|---|---|---|---|
| Untreated, rep 1 | 796.34 | 796.34 | N.D. | N.D. | 9.18 | 7.31 | N.D. | N.D. | 1.76 | 0.92 |
| Untreated, rep 2 | 1445.77 | 1445.77 | N.D. | N.D. | 15.13 | 12.35 | N.D. | N.D. | 1.62 | 0.85 |
| Untreated, rep 3 | 1168.78 | 1168.78 | N.D. | N.D. | 12.46 | 10.08 | N.D. | N.D. | 1.66 | 0.86 |
| 2'-$NH_2$-dCyd, rep 1 | 744.98 | 744.98 | 44.95 | 1.44 | 4.80 | 3.59 | 6.14 | 1.37 | 1.41 | 0.67 |
| 2'-$NH_2$-dCyd, rep 2 | 532.30 | 532.30 | 32.69 | 1.78 | 7.10 | 5.10 | 5.62 | 1.52 | 1.57 | 0.70 |
| 2'-$NH_2$-dCyd, rep 3 | 725.29 | 725.29 | 40.75 | 1.05 | 6.99 | 5.62 | 6.03 | 1.40 | 1.51 | 0.75 |
| 2'-$NH_2$-dUrd, rep 1 | 817.77 | 817.77 | 9.36 | 8.77 | 7.57 | 6.21 | 1.14 | 0.33 | 1.47 | 0.76 |
| 2'-$NH_2$-dUrd, rep 2 | 1007.03 | 1007.03 | 10.65 | 11.73 | 7.82 | 6.70 | 1.06 | 0.42 | 1.26 | 0.66 |
| 2'-$NH_2$-dUrd, rep 3 | 1154.41 | 1154.41 | 12.64 | 12.42 | 10.90 | 8.53 | 1.09 | 0.25 | 1.52 | 0.74 |

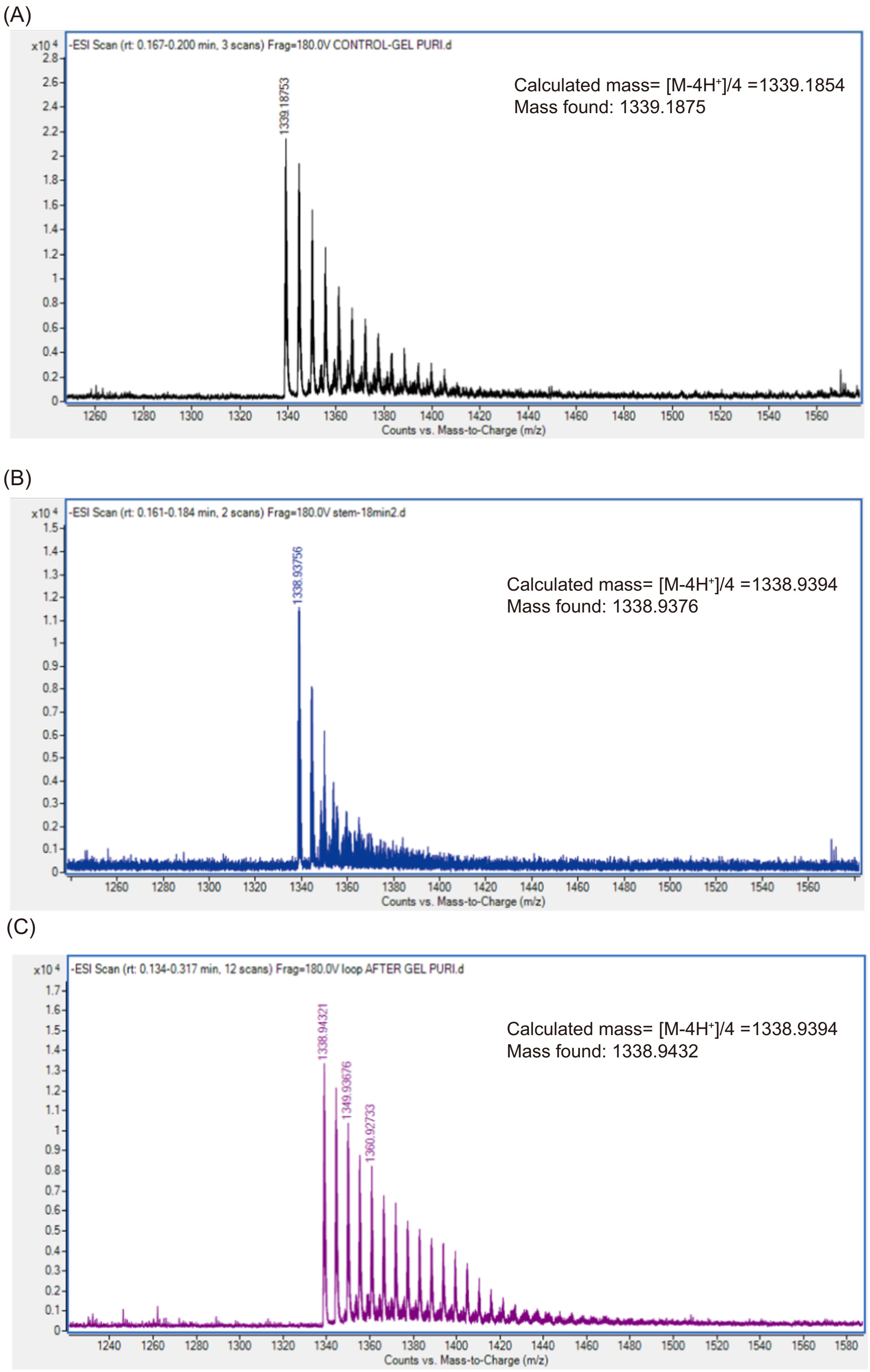


**Supplementary Figure 1.** High resolution mass spectrometry data for synthetic oligos. **(A)** oligo **1**, **(B)** oligo **2**, **(C)** oligo **3**.

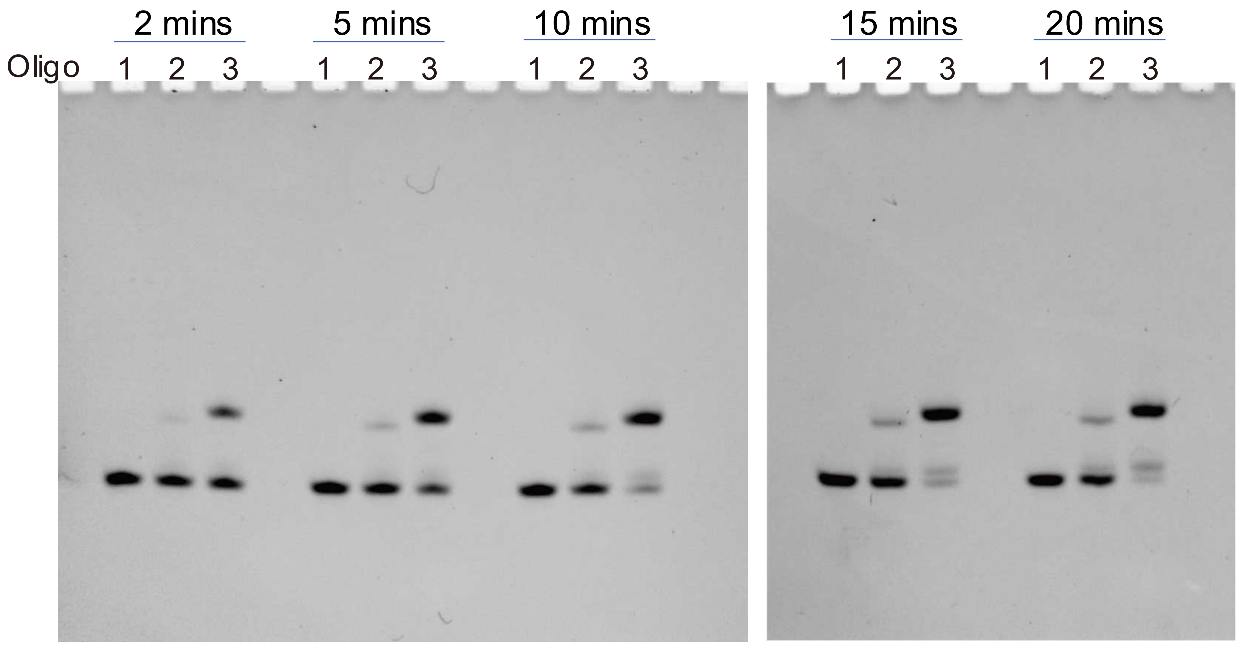


**Supplementary Figure 2.** Time course labeling of synthetic oligos **1-3** with isoS-$N_3$.

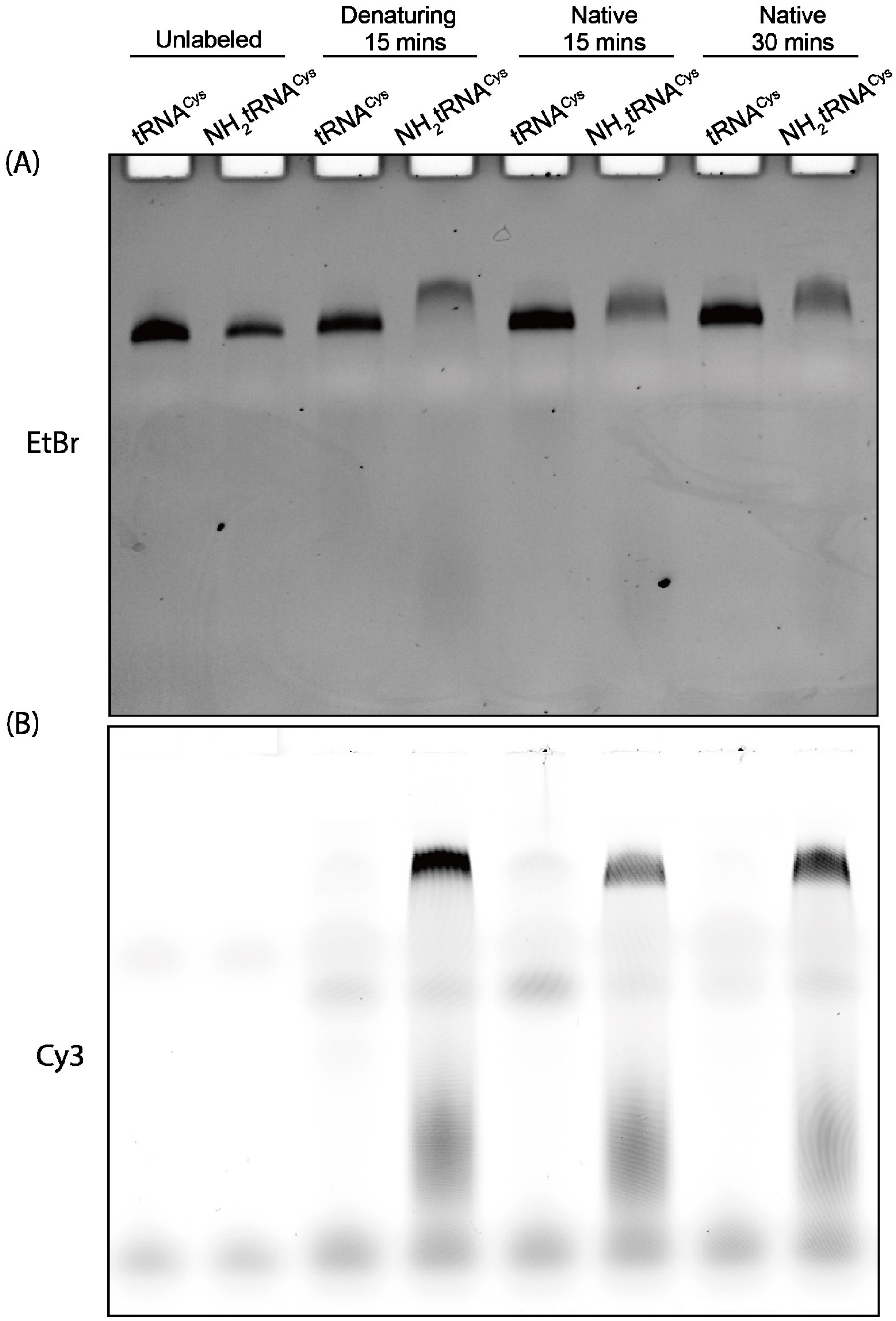


**Supplementary Figure 3.** Reaction of isoS-$N_3$ with 2'-$NH_2$-dCyd-labeled $tRNA^{Cys}$. $tRNA^{Cys}$ or 2'-$NH_2$-C-labeled $tRNA^{Cys}$ were reacted with 5 mM isoS-$N_3$ under denaturing conditions (50% formamide) or folding buffer for the indicated time, and then reacted with an excess of DBCO-Cy3 for 1 hr. The product was detected using EtBr staining **(A)** or by Cy3 fluorescence **(B)**.

A： 500 - 10 ng/uL

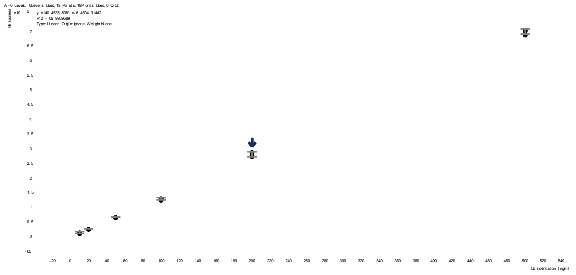

G： 500 - 10 ng/uL

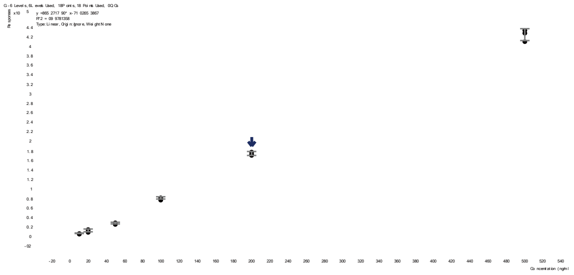

C： 500 - 10 ng/uL

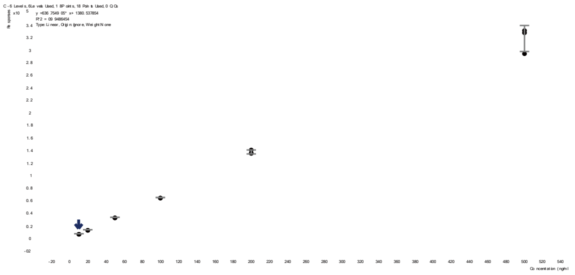

U： 500 - 10 ng/uL

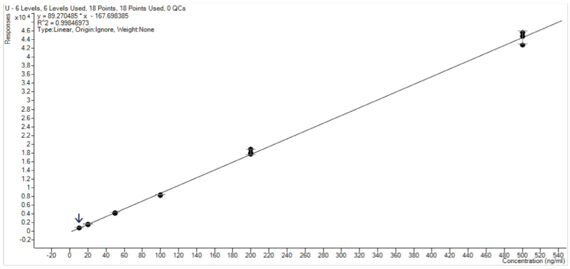


DHU： 50 - 1 ng/uL

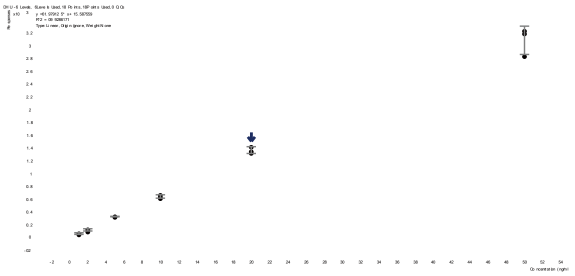

m5C： 50 - 1 ng/uL

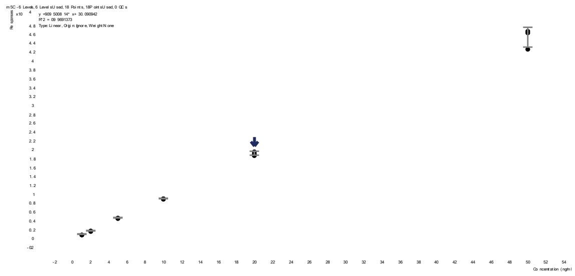

2’-NH2-dCyd： 50 - 1 ng/uL

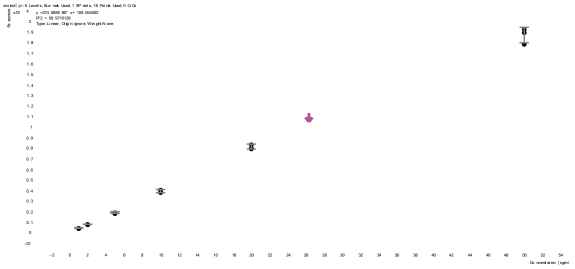

2’-NH2-dUrd： 50 - 1ng/uL

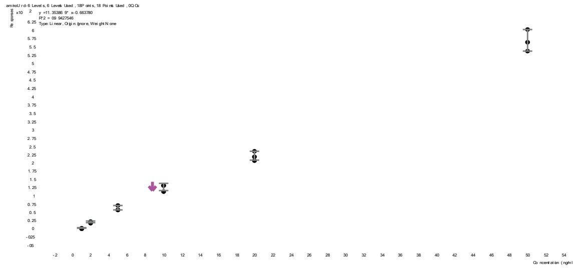

**Supplementary Figure 4**. Representative standard curves for canonical and modified nucleosides analyzed by LC-QQQ-MS.

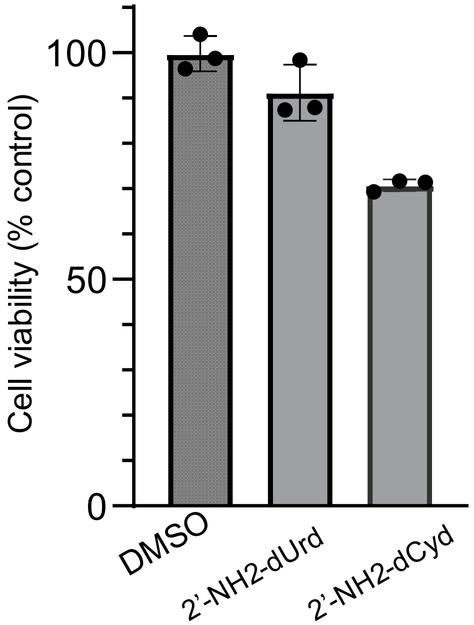


**Supplementary Figure 5.** Cell viability assay after 2'-$NH_2$-dUrd or 2'-$NH_2$-dCyd treatment. HEK293T cells were treated with 1 mM 2'-aminodeoxypyrimidine nucleosides for 18 hr and cell viability was measured using an MTS-based assay. Data represent the mean +/- s.d. (n=3).

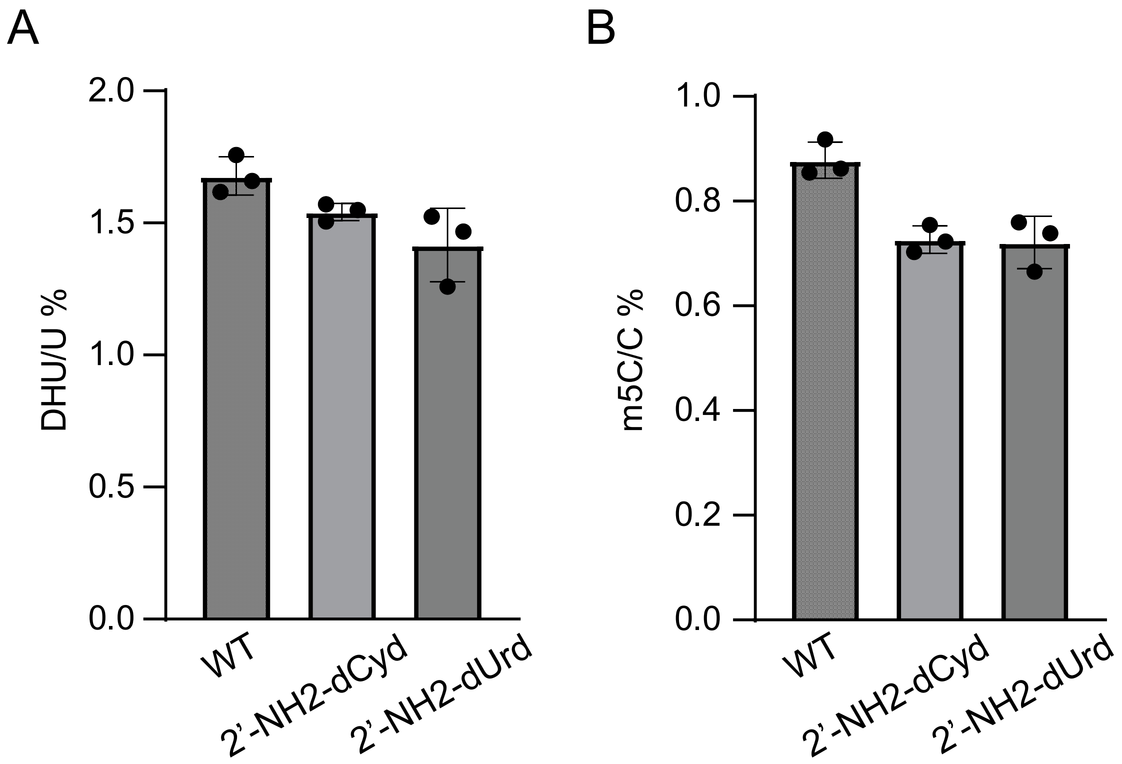


**Supplementary Figure 6.** Levels of endogenous modified nucleotides in RNA isolated from cells treated with 2'-aminodeoxypyrimidine nucleosides. HEK293T cells were treated with 1 mM 2'-$NH_2$-dCyd/dUrd overnight, total RNA was extracted and digested, and D (**A**) and $m^5C$ (**B**) levels were measured by LC-QQQ-MS.

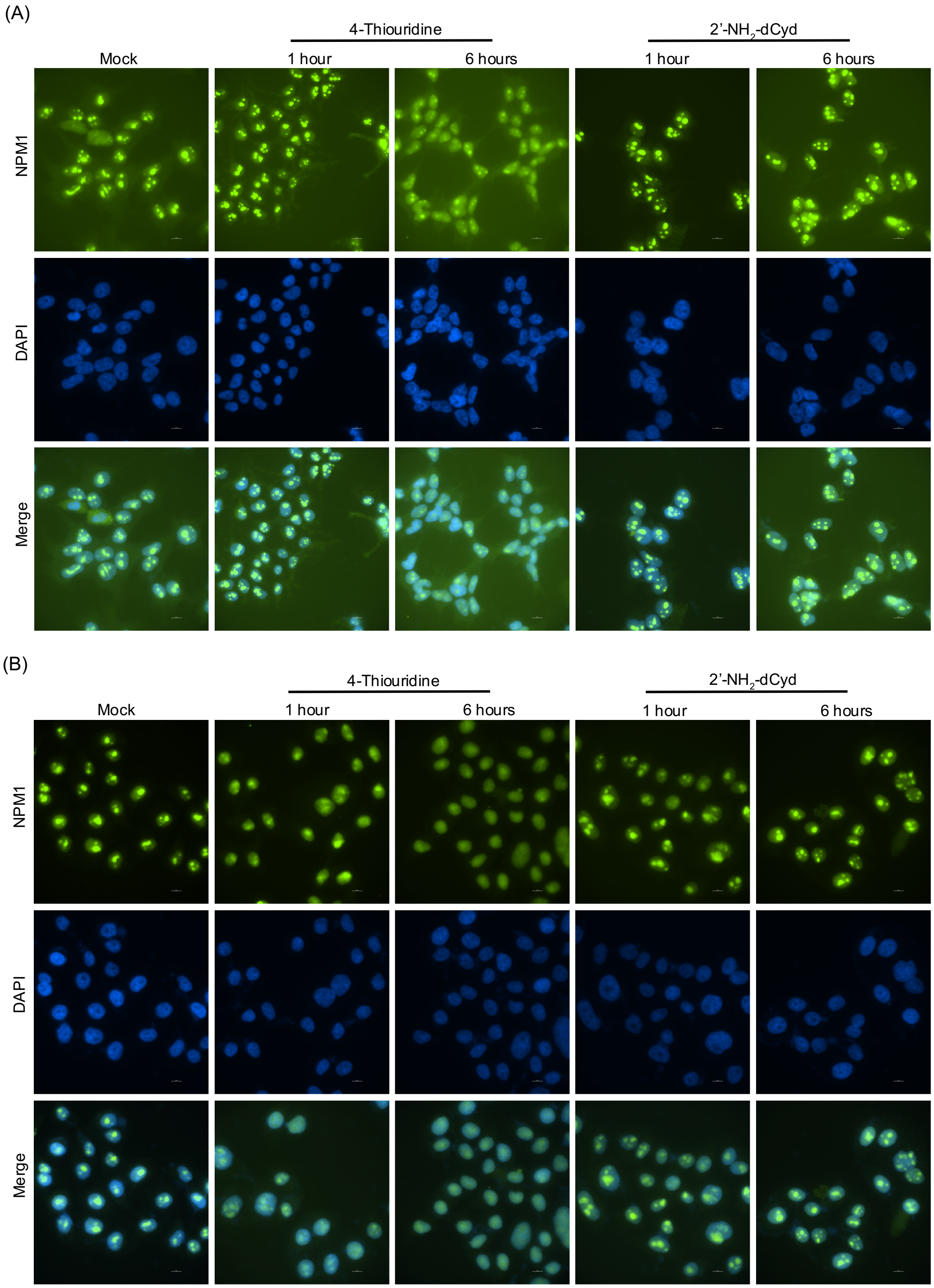


**Supplementary Figure 7**. Analysis of nucleolar structure after treatment with modified pyrimidines. HeLa (**A**) or HEK293T (**B**) cells were treated with 200 μM 4-thiouridine or 1 mM 2'-$NH_2$-dCyd for the indicated time. NPM1 localization was analyzed by immunofluorescence analysis and nuclei were stained with DAPI. Representative images from two independent replicates.

Total RNA *in vitro* labeling with electrophile probes

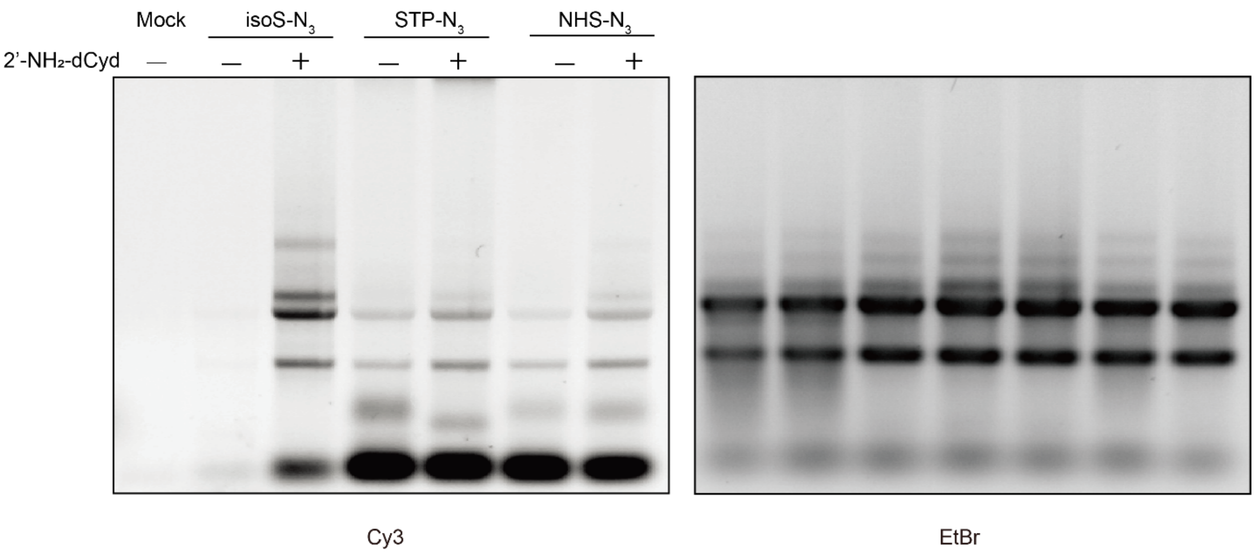


**Supplementary Figure 8.** Labeling of total cellular RNA with different azide-containing electrophiles. RNA was extracted from untreated HEK 293T cells or 1 mM 2'-$NH_2$-dCyd treated HEK293T cells. RNA was labeled with 20 mM NHS-$N_3$, 20 mM STP-$N_3$, or 5 mM isoS-$N_3$ at 37 °C for 15 min in 20 µL folding buffer and then reacted with DBCO-Cy3. Labeled RNA was detected by in-gel fluorescence (left) and EtBr stain (right) as a loading control.

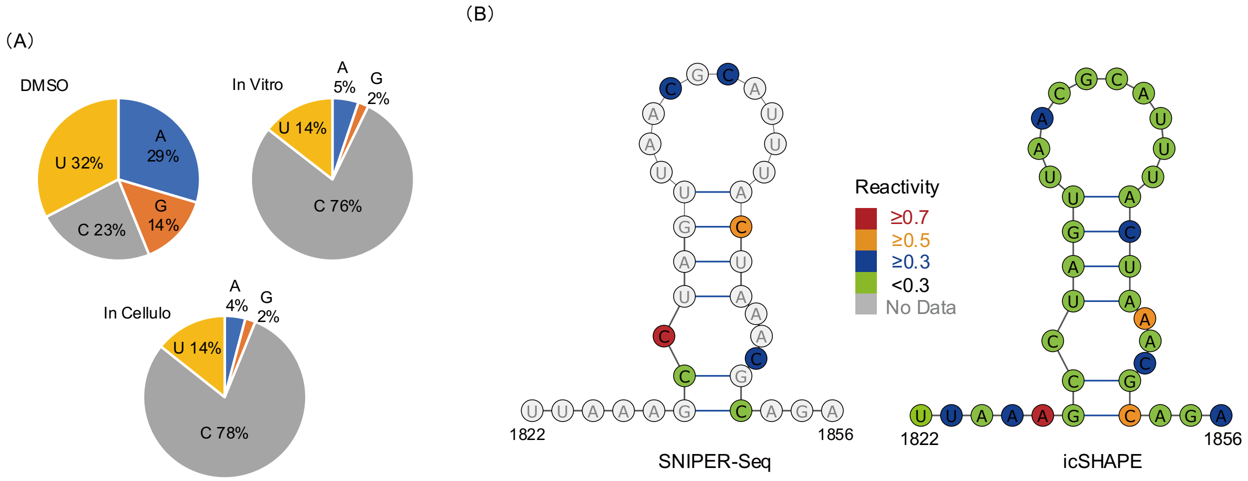


**Supplementary Figure 9.** Transcriptome-wide SNIPER-seq structure probing experiment. **(A)** Transcriptome-wide RT stop distribution from SNIPER-seq samples. Labeling with isoS-$N_3$ was performed on purified 2'-$NH_2$-dCyd-labeled cellular RNA for the *in vitro* sample and performed on cells for the *in cellulo* sample. The DMSO sample was not treated with 2'-$NH_2$-dCyd or isoS-$N_3$. **(B)** Predicted human Malat1 lncRNA (nucleotides 1822-1856) secondary structure with both SNIPER-Seq and icSHAPE reactivity data shown in color from the *in cellulo* sample.

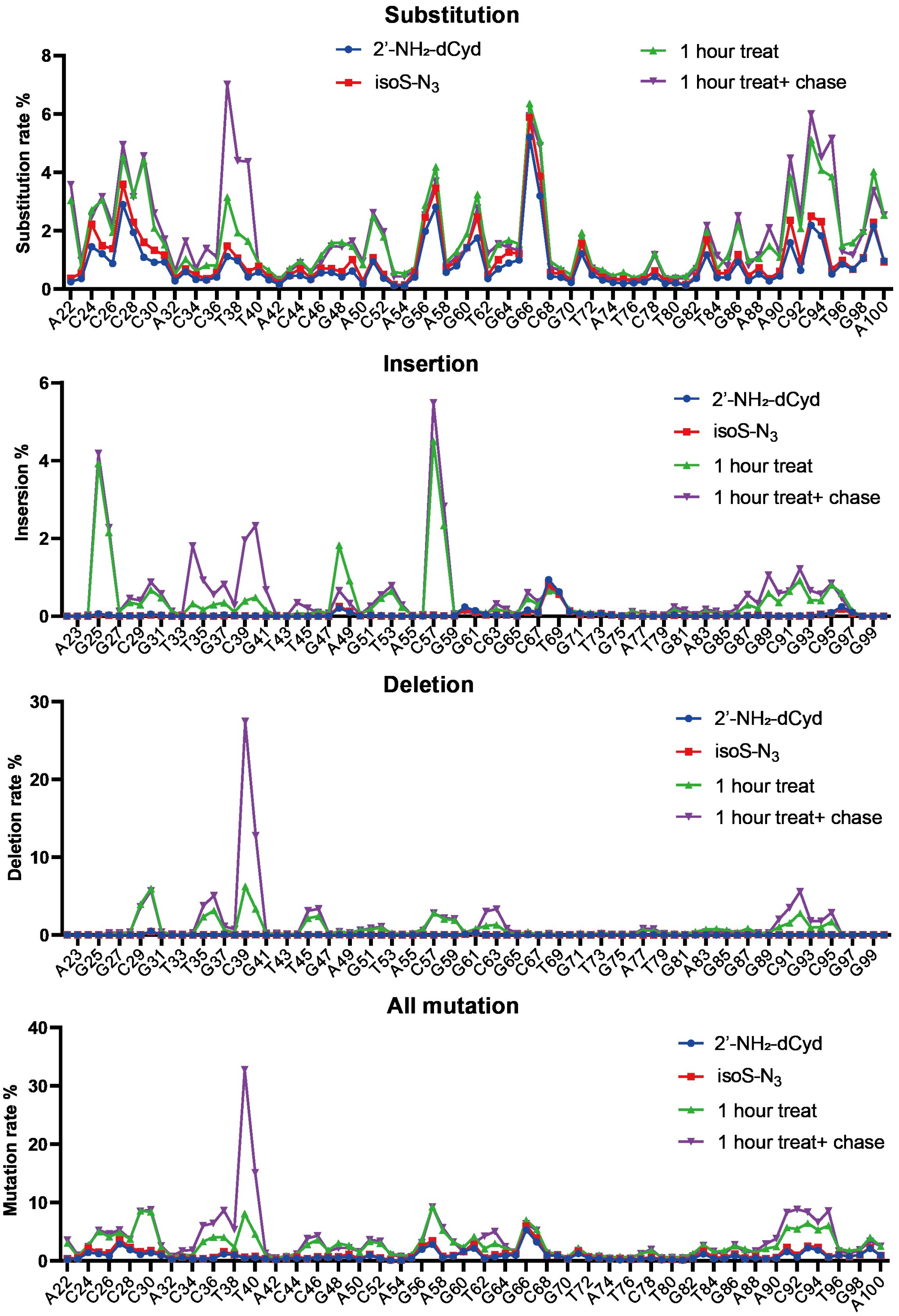


**Supplementary Figure 10.** Mutational profiles of 5S rRNA from SNIPER-seq pulse-chase analysis. HEK293T cells were treated with 1 mM 2’-$NH_2$-dCyd as described, and untreated cells (no 2’-$NH_2$-dCyd feeding or no isoS-$N_3$ treatment) were used as control. Labeled RNA was purified and enriched with DBCO-disulfide-biotin before RT-PCR using gene-specific primers (Supplementary Table 1), amplicon sequencing, and measurement of mutational frequencies.

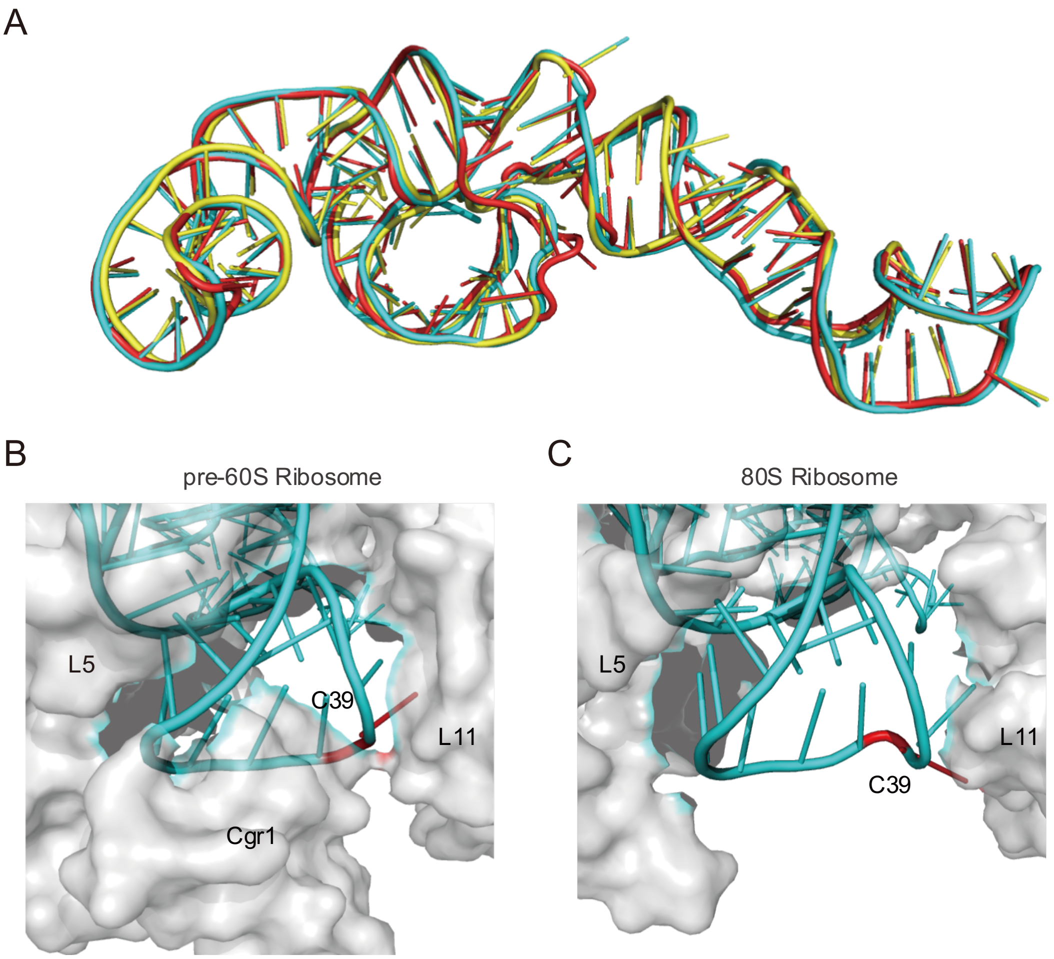


**Supplementary Figure 11**. **(A)** Overlay of 5S rRNA structures from human 80S ribosome (PDB ID: 4UG0, shown in yellow), yeast pre-60S ribosome (PDB: 3JCT, shown in cyan) and yeast 80S ribosome (PDB ID: 4V7R, shown in red). **(B)** Structure of yeast 5S rRNA Loop C (cyan) in complex with ribosomal proteins (gray) from pre-60S ribosome (PDB ID: 3JCT). Residue C39 is colored red. **(C)** Structure of yeast 5S rRNA Loop C (cyan) in complex with ribosomal proteins (gray) from mature 80S ribosome (PDB ID: 4V7R). Residue C39 is colored red.

**Supplementary Scheme 1.** Synthesis of azide-modified electrophile probes.

1).

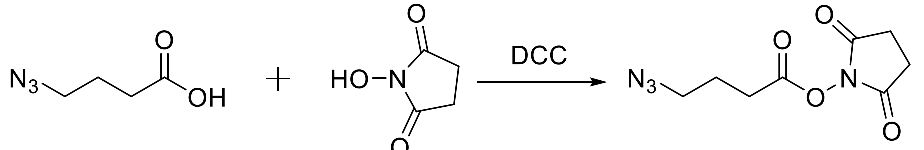


NHS-$N_3$

2).

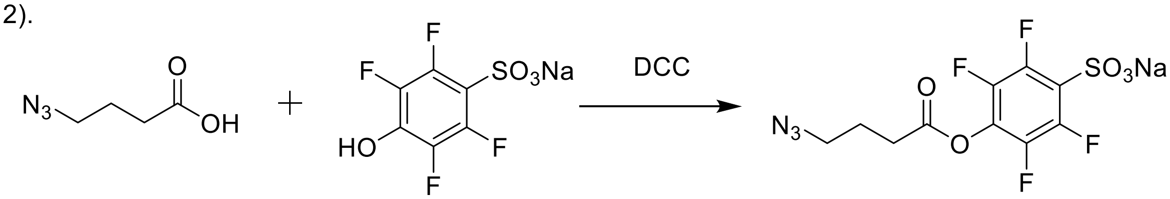


STP-$N_3$

3).

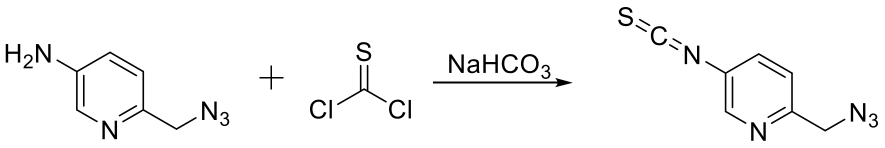


isoS-$N_3$

**NHS-$N_3$.** Synthesis was performed following a published synthetic route[1].

**STP-$N_3$.** To azidobutyric acid (65 mg, 1 mmol) and 4-sulfotetrafluorophenol sodium salt (133 mg, 0.5 mmol) in 1 mL DMF and 15 mL acetone was added 1,3-dicyclohexylcarbodiimide (115 mg, 0.55 mmol) and the mixture was stirred at room temperature for 20 hr. The resulting precipitate was removed by filtration and the filtrate was concentrated under reduced pressure to give a crude product. The product was purified by column chromatography on silica gel with 40% chloroform in acetone as eluant to give 265 mg (71%). $^1$H NMR (500 MHz, DMSO-d6) δ 3.46 (t, J = 6.8 Hz, 2H), 2.87 (t, J = 7.3 Hz, 2H), 1.93 (t, J = 7.1 Hz, 2H).

**isoS-$N_3$.** 60 mg 6-(azidomethyl)pyridin-3-amine[2] was dissolved in 2 mL $CHCl_3$ and an equal volume of saturated aqueous $NaHCO_3$ was added at room temperature. The resulting solution was stirred and 1.0 mL (54 mg, 0.48 mmol) of thiophosgene in 1 mL of $CHCl_3$ was added dropwise. After 1.5 hr, the reaction mixture was filtered and the aqueous layer was extracted twice with 25 mL portions of $CHCl_3$. The combined organic material was dried over anhydrous $MgSO_4$ and concentrated under reduced pressure with a rotary evaporator. The crude product was further purified by column chromatography on silica gel with 20% ethyl acetate in hexane to obtain 50 mg of final product (65%). $^{1}H$ NMR (500 MHz, $CDCl_3$) δ 8.43 (d, J = 2.5 Hz, 1H), 7.48 (dd, J = 8.4, 2.5 Hz, 1H), 7.29 (d, J = 8.3 Hz, 1H), 4.43 (s, 2H). $^{13}C$ NMR (126 MHz, $CDCl_3$) δ 154.00, 146.76, 133.28, 128.89, 122.37, 55.10.

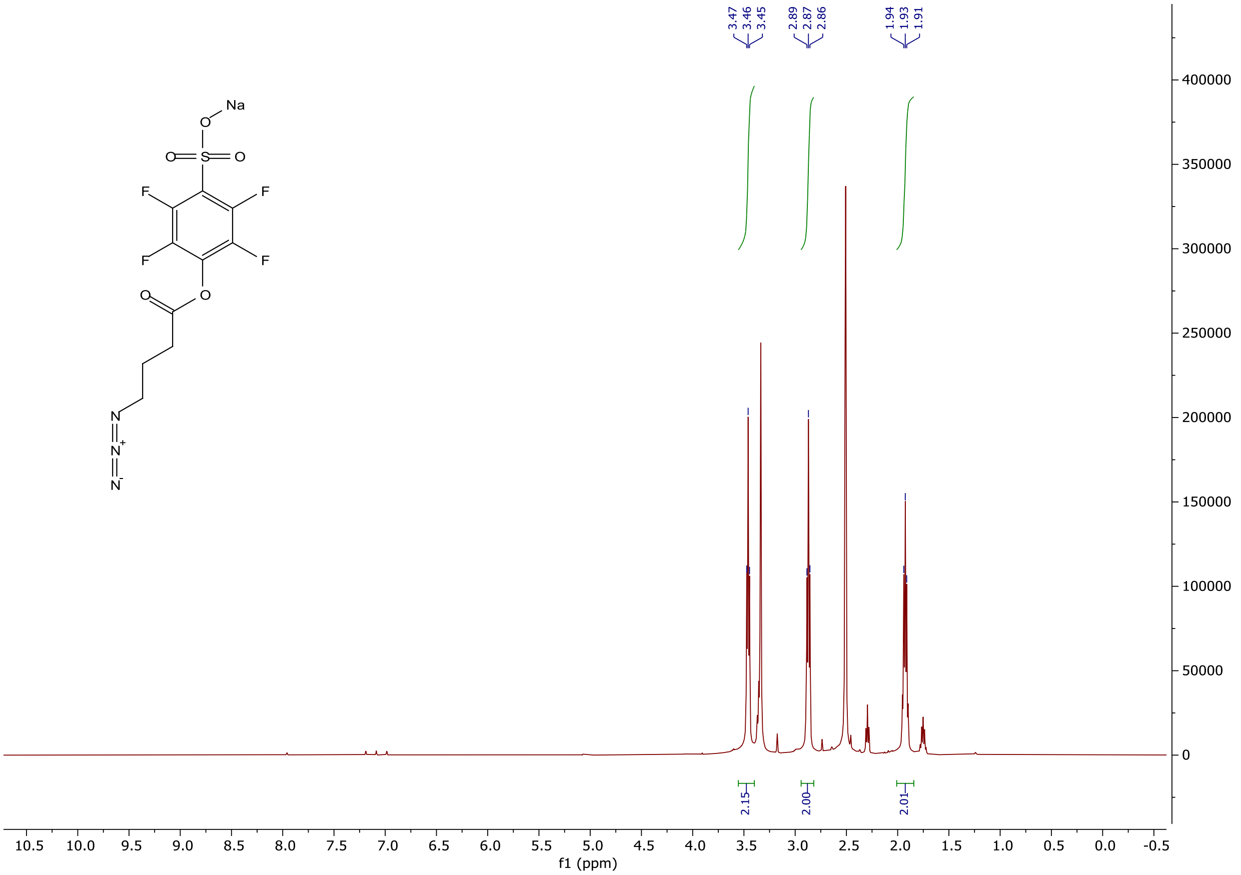

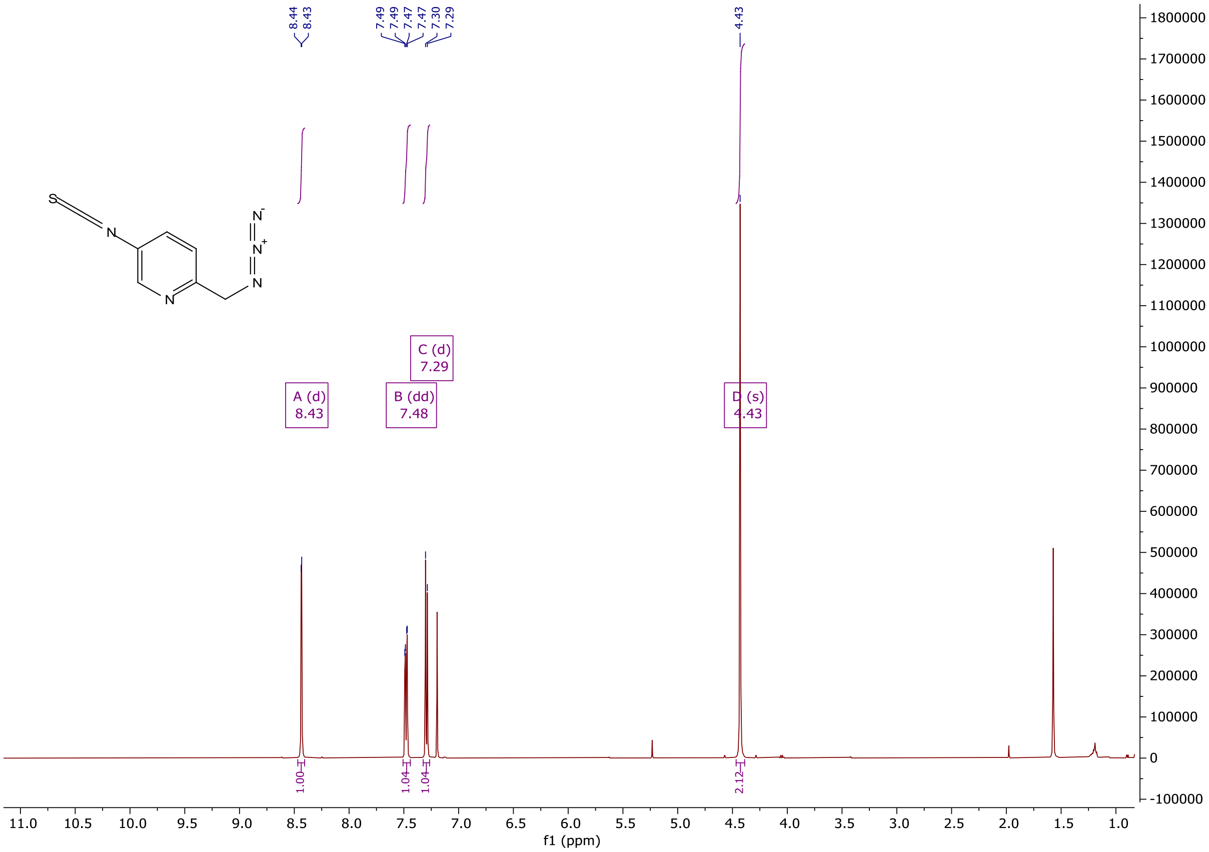
8.44
8.43
7.49
7.49
7.47
7.47
7.30
7.29
4.43
A (d)
8.43
B (dd)
7.48
C (d)
7.29
D (s)
4.43
1.00
1.04
1.04
2.12
f1 (ppm)

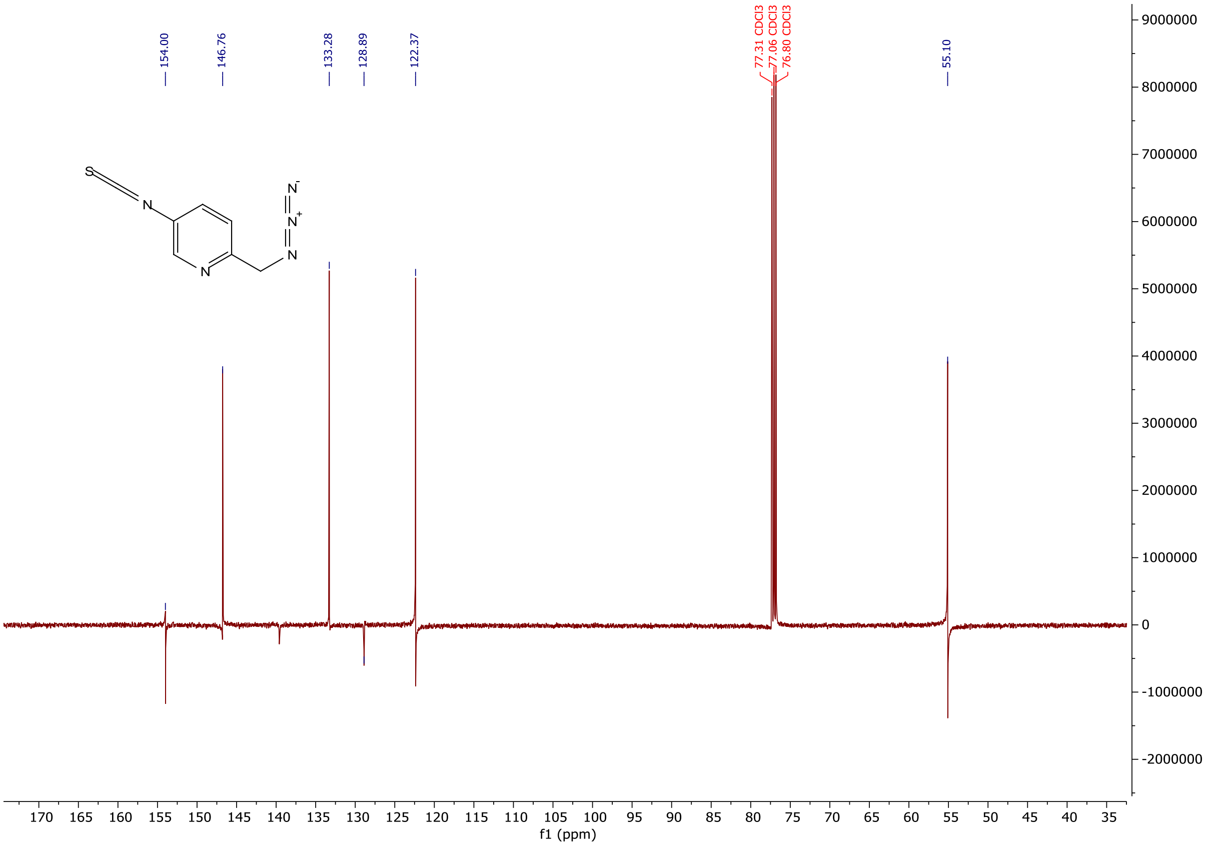
154.00
146.76
133.28
128.89
122.37
77.31 CDCl3
77.06 CDCl3
76.80 CDCl3
55.10
f1 (ppm)
